# Synthesis and Assembly of a New-generation Bifunctional Lipid Nanoparticle for Selective Delivery of mRNA to Antigen Presenting Cells

**DOI:** 10.1101/2023.12.26.572282

**Authors:** Chen-Yo Fan, Szu-Wen Wang, Chi-Huey Wong

**Affiliations:** Genomics Research Center, Academia Sinica, 128 Academia Road, Sec 2, Taipei 115, Taiwan; Department of Chemistry, The Scripps Research Institute, 1050 N. Torrey Pines Rd., La Jolla, CA 92037 United States

**Author notes:** **Corresponding author:** Chi-Huey Wong, Department of Chemistry, The Scripps Research Institute.

**Keywords:** Bifunctional nanoparticle, mRNA vaccine, selective delivery, dendritic cell

## Abstract

Described here is the synthesis and assembly of a new-generation mRNA-bifunctional lipid nanoparticle (mRNA-BLNP3) for selective delivery of mRNA to antigen-presenting cells (APCs). Compared to mRNA-BLNP1 and BLNP2 (BioRxiv,doi.org/10.1101/2023.12.26.572282), the mRNA-BLNP3, incorporating a glycolipid with the lipid moiety designed to target the CD1d receptor on dendritic cells (DCs) and the sugar head group targeting the mannose-binding receptors on APCs is more selective with stronger target-specific immune responses. It was shown that vaccination in mice with mRNA-BLNP1 elicited enhanced cytokine induction and antibody responses compared to traditional mRNA-LNPs, and the mRNA-BLNP2 vaccine, incorporating a mannose-glycolipid, further improved DC targeting. However, BLNP3, incorporating the glycolipid with an aryl-mannose head group targeting the mannose receptor on APCs and the same lipid moiety targeting the Cd1d receptor on DCs, showed superior lymph node targeting *in vivo* with reduced liver accumulation, enhanced mRNA expression in DCs and macrophages, and increased DC maturation. Immunization in mice with mRNA-BLNP3 elicited enhanced humoral and cellular immune responses compared to mRNA-BLNP1 and mRNA-BLNP2, with higher antigen-specific IgG titers and granzyme B–producing CD8^+^ T cells, demonstrating that BLNP3 is a promising bifunctional lipid nanoparticle for delivery of mRNA vaccines with improved efficacy and safety.

## Introduction

While many vaccine candidates against SARS-CoV-2 have been investigated in the past years (1), the speedy development of effective mRNA vaccines demonstrated the most powerful approach to control the pandemic.(2-5) The success of mRNA vaccines depends on the fast design of stable mRNA with nucleotide modification and the subsequent manufacture and encapsulation in lipid nanoparticles (LNPs) for vaccination. Current mRNA vaccines (4, 5) encoding the full length spike protein, contains pseudo-uridine and two proline substitutions to reduce immunogenicity and stabilize the prefusion state, respectively and is formulated with a mixture of four lipids, including a ionizable lipid, a phospholipid, a cholesterol and a PEG-conjugated lipid to form an mRNA-LNP for immunization (6). Vaccination with mRNA is attractive because the mRNA can rapidly translate to the corresponding protein antigen in the cytoplasm after immunization (7, 8) and provide adjuvant activities via toll-like receptors (TLRs) to enhance immune response (7). The success of mRNA vaccines against SARS-CoV-2 has stimulated interests in developing new mRNA vaccines against other diseases and new methods for targeted delivery of mRNA (9) to improve efficacy and safety (10-13).

Among the mRNA delivery systems, non-viral vectors with low immunogenicity represent a simpler and safer alternative to viral vectors. With the advances in development of new biocompatible materials and innovative fabrication approaches, non-viral vectors have become the preferred vehicles for mRNA delivery, and a variety of non-viral vectors have been explored, including lipids and polymers (14, 15). Recent development of LNPs has been focused on encapsulation of mRNA to reduce its degradation by nucleases and provide better transfection efficiency (16) and selective delivery to target specific subsets of antigen-presenting cells (APCs) (17). However, the issue of selectivity remains a challenge (18). Therefore, it is important to develop new LNPs with better selectivity for APCs to enhance immune responses and minimize undesirable translation of mRNA in different cells (19-20). We have previously developed a bifunctional LNP (BLNP1) containing the glycolipid C34 as a selective delivery component to target the CD1d receptor on dendritic cells (DCs) and the T cell receptor on iNKT cells to optimize the Th1 and Th2 pathways in CD4+ and CD8+ T cell responses (21). In addition, we modified the glycan moiety of C34 (22, 23) with mannose to generate BLNP2 with improved selectivity. Herein, we have developed an aryl-mannose glycolipid to target CD1d and DC-SIGN on dendritic cell (DC) and combined with other lipids to generate mRNA-BLNP3 for selective delivery of mRNA to APCs. The spike-mRNA of wild-type SARS-CoV-2 formulated as mRNA-BLNP3 vaccine was shown to elicit stronger immune responses with higher levels of spike protein-specific IgG antibodies than the traditional mRNA-LNP and other mRNA-BLNP vaccines.

## Results

In our first approach, we reduced the four-component LNP commonly used in mRNA formulation to a two-component lipid nanoparticle to simplify the formulation process. We first prepared a guanidinyl lipid and a zwitterionic lipid to form mRNA-GZ-LNP (Fig. 1). In general, LNPs include a helper lipid and a cationic lipid. The helper lipid was used to stabilize the mRNA-LNP delivery system, such as 1,2-dioctadecanoyl-sn-glycero-3-phosphocholine (DSPC)(24) or 1,2-dioleoyl-sn-glycerol-3-phosphoethanolamine (DOPE)(25). These helper lipids not only stabilize the particles but also enhance mRNA delivery efficiency (26). Ionizable lipids are of great importance for delivery of mRNAs because their positively charged head groups can interact with the negatively charged phosphate of the mRNA molecule (27), and protonation of the ionizable components destabilizes the anionic membrane and facilitates nanoparticle disassembly, leading to the release of mRNA to the cytosol (28). To facilitate the mRNA encapsulation and delivery, the ionizable lipid is the most abundant component in LNP formulation. However, the phosphate groups with negative charge in helper lipids allow selective delivery to the spleen for protein expression in spleen and lymph nodes (29). Therefore, we introduce a higher number of zwitterion groups into the formulation of mRNA-LNPs to ensure a more precise mRNA delivery. By connecting a guanidine head with a tertiary amine and a phosphate group in one molecule (L5), the guanidine group is positively charged at physiological and acidic conditions, and the phosphate group is negatively charged at physiological conditions and neutral at acidic conditions. The combination of these functional groups increased the overall phosphate ingredient on lipid nanoparticles (IZ-LNP) to facilitate targeting.

**Figure 1.**
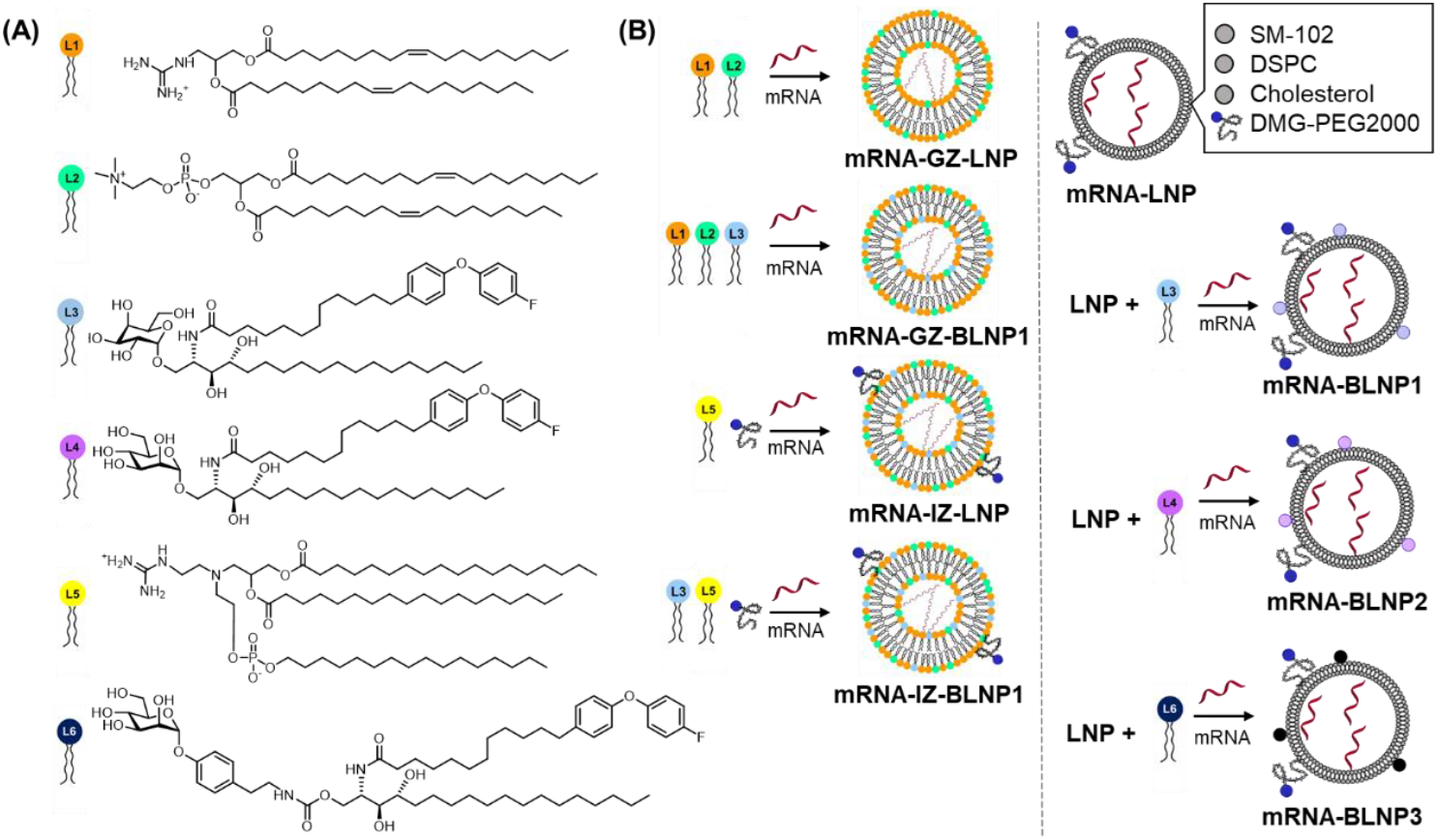
(A) Structure of synthetic lipids, (B) constructs of different mRNA-LNPs and mRNA-BLNPs preparations with bifunctional glycolipid for APC-targeted delivery

To improve target delivery, we then prepared different bifunctional glycolipids with the sugar head targeting APCs and the lipid moiety targeting CD1d on DCs (Fig. 2) and include the bifunctional lipid in the mRNA-GZ-BLNP, mRNA-IZ-BLNP and mRNA-BLNP formulations. Using these BLNPs, we can synthesize nanoparticles in sizes of one to two hundred nm with high transfection efficiency in HEK293 cells (BioRxiv, doi.org/10.1101/2023.12.26.572282). Although it is practical to use these BLNP formulation systems for *in vitro* or *in vivo* study, the long-term storage and serum instability of these BLNPs was still a concern due to the absence of cholesterol and PEGylated lipids. When mRNA-GZ-BLNP1 was incubated with 10% FBS at 37 °C for 6-48 h, we found a gradual increase of particle size as the incubation time increased (BioRxiv, doi.org/10.1101/2023.12.26.572282). Interestingly, inclusion of L3 showed a slightly stabilization of LNP, while inclusion of cholesterol and PEGylated lipid improved the transformation of LNPs to multilayer nanoparticles. Therefore, for *in vivo* studies, 1.5% PEG lipid and 38.5% cholesterol were included in mRNA-GZ-BLNP1 or mRNA-IZ-BLNP1 to improve stability. In the assembly of mRNA with GZ-LNP/IZ-LNP and mRNA mixing with L3, we observed an average particle size of mRNA-GZ-BLNP1 or mRNA-IZ-BLNP1 at around 174 or 140 nm by DLS, respectively with spherical morphology revealed by TEM and CryoEM (BioRxiv, doi.org/10.1101/2023.12.26.572282). We then immunized mice with spike-encoding mRNA-LNPs and a significantly higher level of neutralizing antibodies in the mRNA-GZ-BLNP1 group was found. In addition, the IgG titer in mice immunized with mRNA-GZ-BLNP1 of WT, Delta, or Omicron spike protein was determined, and it was found that the WT mRNA-BLNP1 vaccine elicited high IgG titers against the WT spike protein, but the titer was reduced against the Delta and Omicron spike proteins. However, though the WT mRNA vaccine with deletion of glycosites in the stem (WT-deg-CD/HR2)-GZ-BLNP1) induced a similar level of IgG titers against the WT spike protein, it generated significantly higher IgG titers against the Delta and Omicron spike proteins (30). On the other hand, when we incorporated the bifunctional lipid L3 into a conventional LNP formulation (Fig. 1B), we observed a strong induction of both cytokines IL-4 and IFN-γ (BioRxiv, doi.org/10.1101/2023.12.26.572282) and a selective uptake of mRNA-BLNP1 by BMDCs but not by the non-BMDCs cells in flow cytometry analysis. The other mRNA-BLNP2 with mannose as the head group also showed a similar phenomenon, indicating that the BMDC uptake was mediated by the lipid moiety. In a separate study, the formulated mRNA–LNPs were incubated with dendritic cells (DCs), B cells, and T cells to evaluate cell-type specificity. Both mRNA-BLNP1 and mRNA-BLNP2 exhibited a pronounced preference for DCs compared with B cells and T cells. Notably, mRNA-BLNP2 demonstrated the highest DC targeting efficiency, with 38.3% FITC-positive cells, which is likely attributed to dual targeting mediated by CD1d and mannose-binding receptors on DCs.

**Figure 2.**
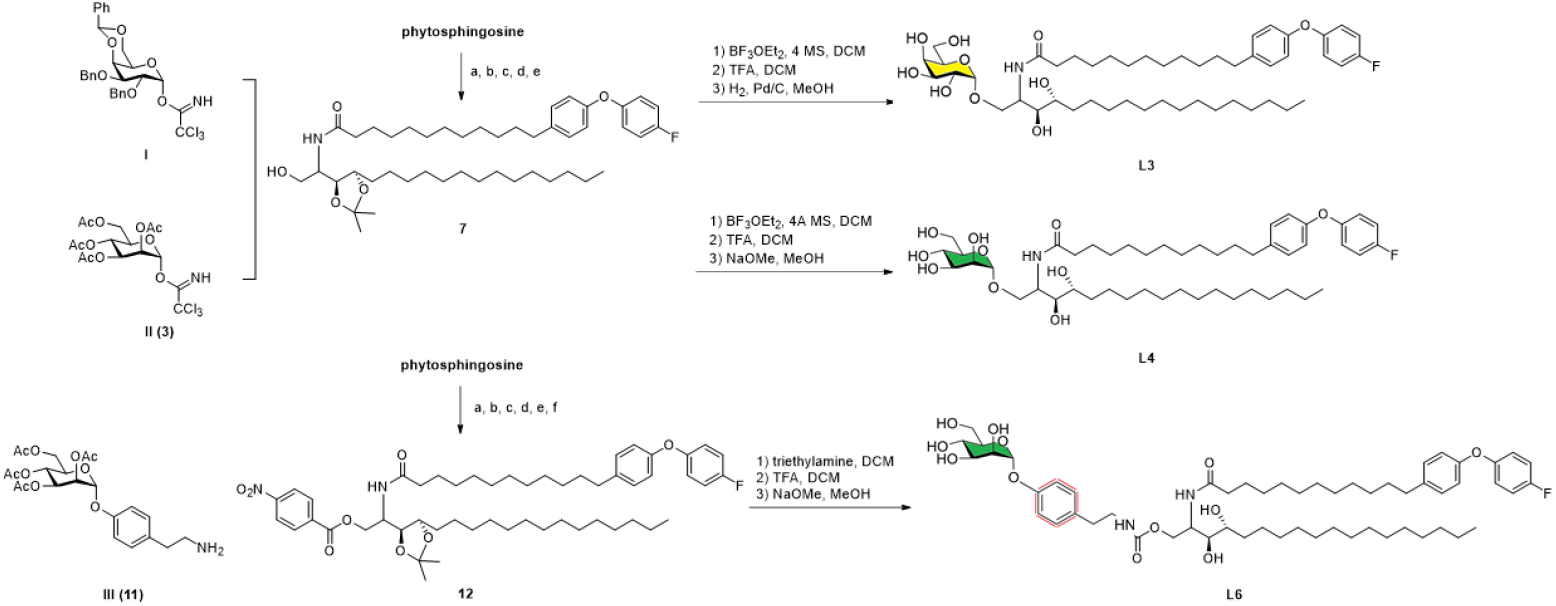
Preparation of glycolipid **L3, L4** and **L6**. Reagents and conditions: (a) 11-bromoundecanoic acid, PPh_3_, toluene, reflux, 24h, 76%; (b) 4-(4-fluorophenoxy)benzaldehyde, LHMDS, THF, 0°C to rt, 12h, 69%; (c) H_2_, Pd/C, MeOH, 2h, quant; (d) phytosphingosine, HBTU, DIPEA, THF, 12h, 80%; (e) 2,2-DMP, PTSA, 2h, quant; (f) 4-nitrophenyl chloroformate, Et_3_N, THF, 12h. Compounds **I**(22), **II**(31), **III**(31) were synthesized according to the reference respectively.

With this promising result, we decided to incorporate L6 (Fig. 2), which contains the aryl-mannose head as a better ligand for DC-SIGN (31), into the design of new-generation BLNP (BLNP3). We next evaluated whether the mRNA delivery specificity and enhanced innate immune responses observed with CD1d- and mannose-receptor-targeted BLNPs *in vitro* could be recapitulated *in vivo*. C57BL/6 mice were intramuscularly (i.m.) immunized in the leg muscle with luciferase (Luc) mRNA-loaded BLNP3 or conventional LNP, and luciferase expression was monitored by in vivo bioluminescence imaging at 1, 6, and 24 h post-injection (Fig. 3A). Both formulations primarily transfected the injection site and, to a lesser extent, other organs. Because restricting mRNA transfection to the administration site and draining lymph nodes are critical to minimizing systemic toxicity, we analyzed the biodistribution of LNPs in major organs and inguinal lymph nodes (iLNs). The *Ex vivo* luminescence imaging revealed no significant accumulation in the heart, spleen, lung, or kidney for either BLNP3 or conventional LNP, with signal detected mainly in the liver and iLNs (Fig. 3B). However, BLNP3 administration resulted in a marked reduction of hepatic radiance compared with conventional LNP, where liver signal accounted for 75% of total intensity, confirming the superior lymph node-targeting capacity of BLNP3 (Fig. 3C).To investigate which cell types in the lymph node were transfected by LNPs, eGFP-encoding mRNA was delivered and detected by flow cytometry. It was found that BLNP3 successfully delivered eGFP mRNA to antigen-presenting cells, including macrophages and dendritic cells (DCs), with DCs (2.6-fold) and macrophages (1.4-fold) exhibiting higher mRNA expressions compared to LNPs (Fig. 4B and C). Since enhanced expression of mRNA in APCs is important for the activation of adaptive immunity and targeting CD1d or DC-SIGN can stimulate DC maturation (32), we analyzed matured DCs (CD80+CD86+) using flow cytometry and found that mRNA-BLNP3 markedly increased the percentage of matured DCs in lymph node (Fig. 4D and E). In contrast, LNP only modestly induced DC maturation, suggesting that the mRNA-BLNP3 formulation can enhance the selective delivery of mRNA to APCs and activate DCs. All these observations were further supported with cellular uptake assay (Fig. S1) and flow cytometry analysis (Fig. S2-S5).

**Figure 3.**
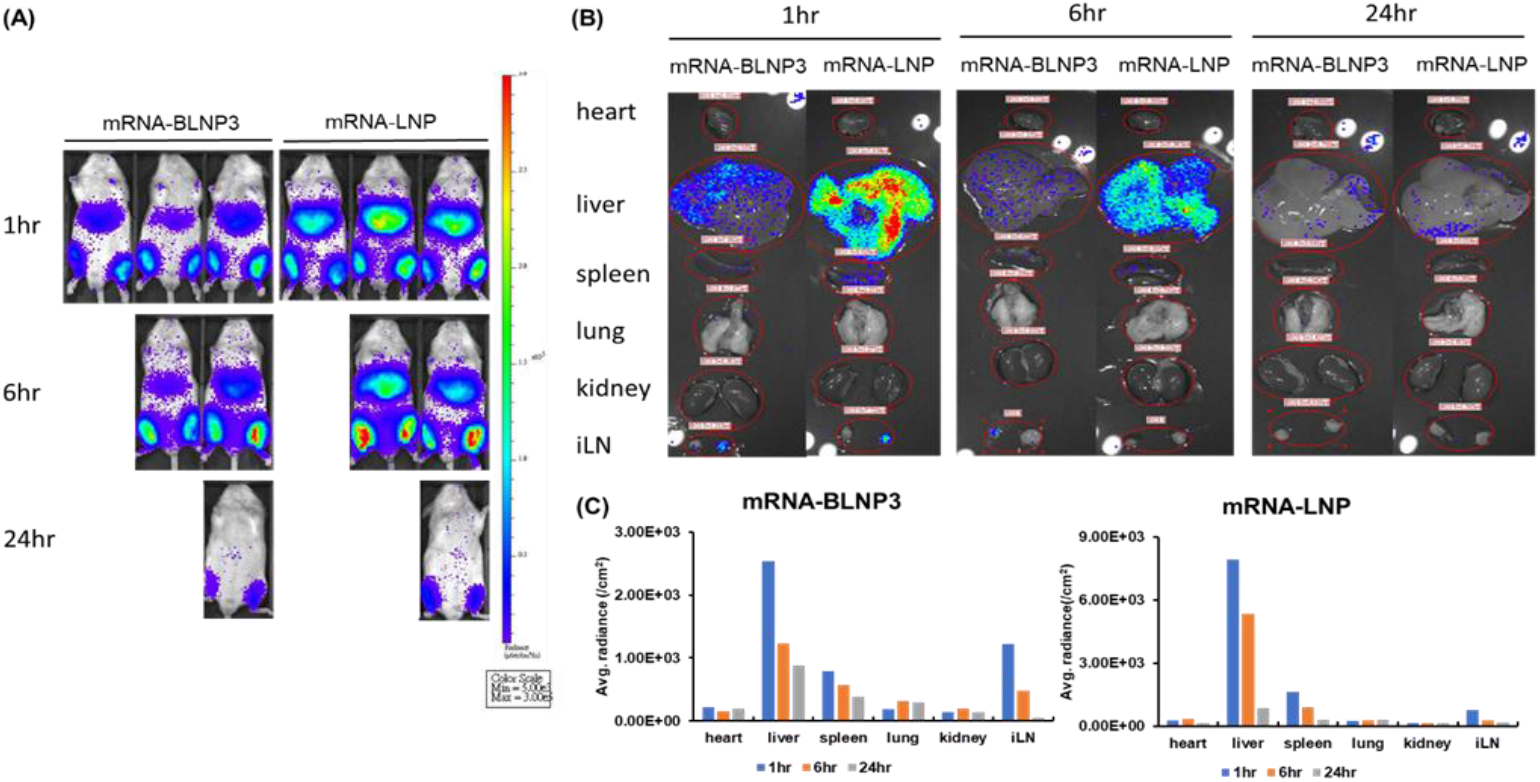
Selective mRNA delivery with mRNA-BLNP3 formulation. (A) *In vivo* bioluminescence imaging at 1, 6 and 24 h post-treatment of Luc mRNA-loaded LNP vs BLNP3. Mice were i.m. injected with Luc mRNA-loaded LNP or Luc mRNA-BLNP3 (5 μg mRNA per mouse). Total luminescence was quantified at different time points. (B) *Ex vivo* luminescence imaging. Mice were i.m. injected with Luc mRNA-LNP or Luc mRNA-BLNP3 (5 μg mRNA per mouse) 1, 6, 24 h before they were sacrificed. Major organs and iLNs were collected for *ex vivo* imaging. (C) Quantification of luminescence signals in B.

**Figure 4.**
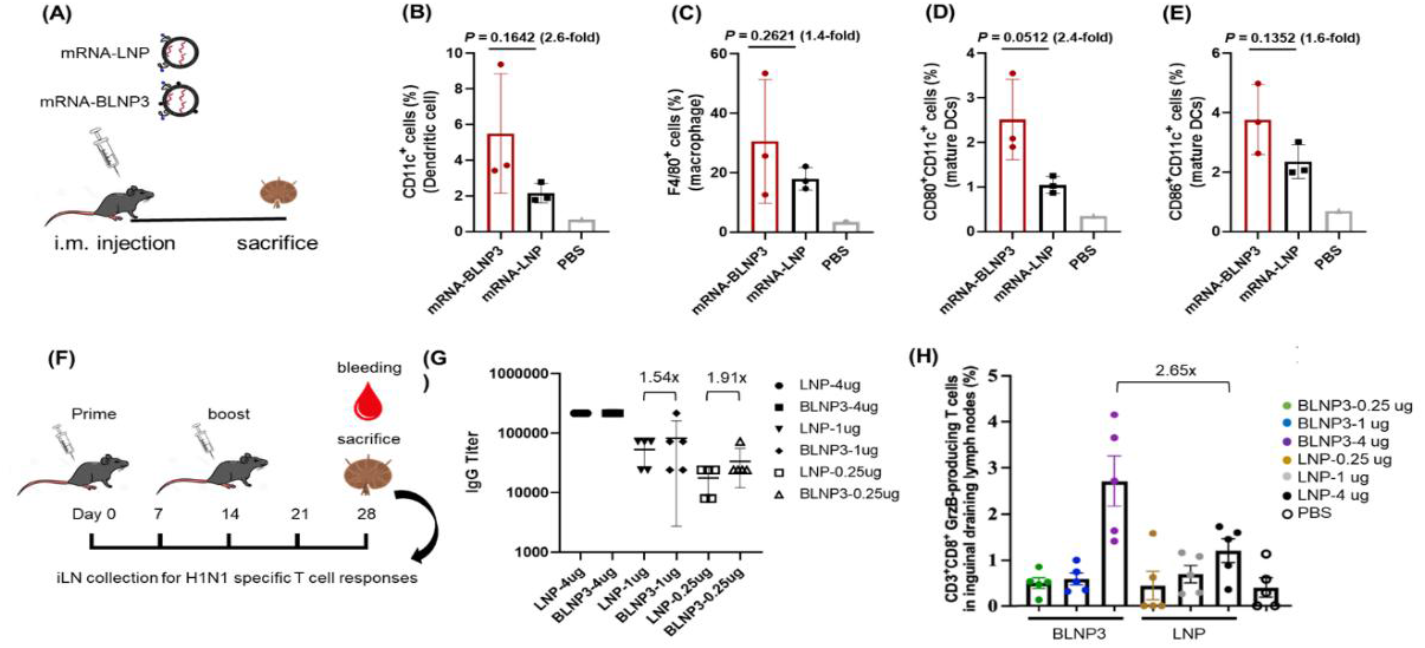
(A) Mice were i.m. injected with eGFP mRNA-LNP and eGFP mRNA-BLNP3 (5 μg mRNA per mouse) 6 h before they were sacrificed, and iLNs were collected for FACS analysis. The error bar around each data point is the SD. Flow cytometry analysis of (B) DCs (C) macrophages, and matured DCs: (D) CD80^+^CD11c^+^ cells and (E) CD86^+^CD11c^+^ cells in iLNs (n = 3). (F) Cellular immune responses induced by H1N1 mRNA LNP vaccine: scheme of the prime, boost vaccination and the analysis of the humoral and T cell responses. C57BL/6J mice were i.m. immunized twice with 4, 1 or 0.25 μg of H1N1-mRNA LNP or H1N1-mRNA BLNP3 vaccine on days 0 and 14. (G) H1N1-specific IgG antibody titers in the mice treated with H1N1-mRNA-LNP or H1N1-mRNA-BLNP3 at different dosages. Analysis of cytotoxic T cell immunity induced by H1N1-mRNA vaccines in mice (H) The percentage of GrzB-producing cells in CD3^+^CD8^+^ T cell populations. Mean ± standard error of the mean (SEM) was plotted, and the statistical significances compared to PBS group were determined by one-way ANOVA with Prism 10.4.1.

Next, we evaluated humoral and T cell responses in immunized mice. Vaccination followed the timeline depicted in Fig. 4F, with a prime dose on day 0 and a boost on day 14. Although no significant difference was observed at the higher dose (4 μg), H1N1-mRNA-BLNP3 elicited higher anti-H1 IgG titers than conventional H1N1-mRNA-LNP at 1 μg and 0.25 μg doses (Fig. 4G). Moreover, immunization with H1N1-mRNA-BLNP3 elicited a 2.65-fold increase in the frequency of granzyme B–producing CD3^+^CD8^+^ T cells compared to H1N1-mRNA-LNP (Fig. 4H). These results indicate that the mRNA-BLNP3 formulation enhances both humoral and cellular adaptive immune responses compared to the clinically relevant mRNA-LNP vaccine. Collectively, these findings demonstrate that the mRNA-BLNP3 formulation can significantly improve the immunogenicity of mRNA vaccines, underscoring their translational potential.

## Discussion

In this study, we compared a series of bifunctional lipid nanoparticle (BLNP) formulations for enhancing the specificity and immunogenicity of mRNA vaccines and simplified the conventional LNP systems into guanidinium/zwitterion-based BLNPs. We successfully achieved selective targeting of antigen-presenting cells, particularly dendritic cells (DCs), via CD1d and mannose-binding receptor pathways. Among them, the mENA-BLNP3 formulation exhibited superior lymph node targeting while minimizing off-target liver accumulation, and experiments *in vivo* demonstrated significantly enhanced mRNA expression in DCs and macrophages, increased DC maturation (CD80^+^CD86^+^), and elevated cytokine responses, including IFN-γ and IL-4. Importantly, the mRNA-BLNP3 formulation also promoted stronger humoral and cellular adaptive immune responses, including higher IgG titers and increased granzyme B–producing CD8^+^ T cells, especially at lower antigen doses. These findings highlight the potential of bifunctional glycolipids for designing next-generation mRNA delivery systems that not only improve cellular targeting but also vaccine efficacy. Taken together, the bifunctional BLNP3 represents a promising candidate for the development of mRNA-BLNP3 vaccines with enhanced immunological performance and safety profiles.

## Materials and Methods

All study and synthesis (Schemes S1 and S2) data are included in the article and/or *<u>SI Appendix</u>*.

## Author Contributions

C.-Y.F. and C.-H.W. designed research, C.-Y.F., S.-W.W. and J.-Y.C. performed data analysis, C.-Y.F. and C.-H.W. prepared draft and final writing.

## Co-Authors

Chen-yo Fan, Genomics Research Center, Academia Sinica. Email: cyfan@rockbiomedical.comSzu-wen Wang, Genomics Research Center, Academia Sinica.

## Acknowledgments

This research was supported by Academia Sinica. We thank CY Chen for assistance in experiment and the National RNAi Core Facility at Academia Sinica for related services.

## Supporting Information

### Materials and Methods

For chemical synthesis, all starting materials and commercially obtained reagents were purchased from Sigma-Aldrich and used as received unless otherwise noted. All reactions were performed in oven-dried glassware under nitrogen atmosphere using dry solvents. ^1^H and ^13^C NMR spectra were recorded on Brucker AV-600 spectrometer and were referenced to the solvent used (CDCl_3_ at δ 7.24 and 77.23, CD_3_OD at δ 3.31 and 49.2, and D_2_O at δ 4.80, and DMSO-d_6_ at δ2.5 and 39.51 for ^1^H and ^13^C, respectively). Chemical shifts (δ) are reported in ppm using the following convention: chemical shift, multiplicity (s = singlet, d = doublet, t = triplet, q = quartet, m = multiplet), integration, and coupling constants (*J*), with *J* reported in Hz. High-resolution mass spectra were recorded under ESI-TOF mass spectroscopy conditions. Silica gel (E, Merck) was used for flash chromatography. Transmission electron microscopy (TEM) images were obtained by a FEI Tecnai G2 F20 S-Twin. EM imaging was conducted in bright-field mode at an operating voltage of 200 kV. Images were recorded at a defocus value of ∼1.8 µm under low-dose exposures (25–30 e/Å2) with a 4k × 4k charge-coupled device camera (Glatan, Pleasanton, CA, USA) at a magnification of 50,000×. Experiments were carried out at the Academia Sinica Cryo-EM Facility (Taipei, Taiwan). mMESSAGE mMACHINE® Kit and SYBR™ Safe DNA Gel Stain is from Thermo Scientific™. 18:1 PE CF and DMG-PEG2000 are from Avanti Polar Lipids, Inc. Cholesterol and DSPC are from Sigma Aldrich. SM-102 is from MedChemExpress. IFNγ and IL4 ELISpot kit are from Mabtech AB.

### Experimental Section

#### Cell Cultures

Human Embryonic Kidney Cells 293 (HEK293T cell) were cultured in Dulbecco’s modified Eagle’s medium (DMEM) (Thermo Scientific™) supplemented with 10% fetal bovine serum (FBS) (Thermo Scientific™) and 1% Gibco™ Antibiotic-Antimycotic (anti-anti) (Thermo Scientific™).

#### Animals and Immunization

Balb/c mice (8 weeks) were purchased from National Laboratory Animal Center, Taiwan. All the mice were maintained in a specific pathogen-free environment. BALB/c mice aged 6 to 8 wk old (n = 5) were immunized intramuscularly with 15 μg of mRNA-LNPs in phosphate-buffered saline (PBS, 100 μL) at wk 0 and boosted with a second vaccination at wk 2, and serum samples were collected from each mouse 1 wk after the second immunization and subjected to ELISA analysis. The experimental protocol was approved by Academia Sinica’s Institutional Animal Care and Utilization Committee (approval no. 22-08-1901).

#### Antibodies and Proteins

SARS-CoV-2 full-length spike protein was purchased from ACROBiosystems. HRP conjugated anti-mouse secondary antibody and horseradish peroxidase substrate were from Thermo Scientific. Mouse monoclonal anti–β-actin were purchased from Millipore. All commercial antibodies were validated for specificity by companies via Western blot.

#### Construction of mRNA synthesis plasmid

The DNA sequence of WT (Wuhan/WH01/2019 strain) SARS-CoV-2 spike (1-1273) with K986P and K987P mutation (2P) were codon-optimized for Homo sapiens. The spike DNA sequence was cloned into pMRNAXP mRNA Synthesis Vector (System Biosciences) that contains T7 promoter, 5’ and 3’ UTR and poly A tail. Briefly, pMRNAXP vector was digested with EcoRI and BamHI (Thermo Scientific™) at 37°C for 1 hour. DNA sequence of the spike protein was amplified by KOD OneTM PCR master mix (TOYOBO Bio-Technology). The linearized pMRNAXP vector and the PCR fragments of DNA encoding spike were cleaned up by Wizard® SV Gel and PCR Clean-Up System (Promega). The PCR fragments of spike protein DNA were cloned into linearized pMRNAXP vector using In-Fusion® HD Cloning Kit (Clontech® Laboratories, Inc.). The cloning mixtures were transformed to One Shot™ TOP10 Chemically Competent E. coli (Thermo Scientific™) and incubated at 37°C overnight.

#### In vitro transcription of mRNA

mRNA was produced using mMESSAGE mMACHINE® Kit (Thermo Scientific™) according to the user guide. Briefly, DNA was linearized by NdeI (Thermo Scientific™) downstream of the poly A tail at 37°C for 1 hour and cleaned up. The linearized DNA was in vitro transcribed using T7 RNA polymerase in the presence of cap analog [m7G(5’)ppp(5’)G] and NTP at 37°C for 2 hours. DNase was added to the mixture and incubated at 37°C for 15 minutes to digest double-stranded DNA. mRNA was precipitated by lithium chloride (LiCl) and the mixture was centrifuged to remove unincorporated nucleotides and protein. The mRNA pellet was washed with 70% ethanol and re-centrifuged to remove remaining unincorporated nucleotides. The mRNA was resuspended in nuclease-free water and stored at - 20°C until use.

#### Fabrication of mRNA-lipid nanoparticles

The composition of lipids including SM-102, DMG-PEG2000, DSPC (1,2-distearoyl-sn-glycero-3-phosphocholine), cholesterol and DMG-PEG2000 were prepared in ethanol at a molar ratio of 50:10:38.5:1.5. mRNA was dissolved in 10 mM pH 4 citric acid buffer. The weight ratio of lipid: mRNA was 3:1 and the nitrogen-to-phosphate (N/P) ratio was 3. For syringe injections, organic phase and aqueous phase were injected via syringes simultaneously. The mixture was then diluted 2 times with PBS. For microfluidic device, organic phase and aqueous phase were passed through the channel on the cartridge using NanoAssemblr® Ignite™ (Precision NanoSystems) and the mixture were then diluted 2 times with PBS. The flow rate ratio (FRR) (aqueous phase and organic phase) was 6. The formed mRNA-LNP was dialyzed against PBS by MWCO 3.5 kDa dialysis membrane (SPECTRUMLABS®.COM)

#### Quantification of encapsulated mRNA

Encapsulation efficiency was determined by Quant-iTTM RiboGreenTM RNA Reagent and Kit (Thermo Scientific™). Samples were diluted 250-fold with 1X TE buffer and further diluted 2-fold with TE buffer or TE buffer containing 2% Triton X-100. Ribosomal mRNAs were prepared as 100, 50, 25, 12.5 and 0 ng/ml in TE or TE buffer containing 1% Triton X-100 to establish the standard curve. After incubation at 37 ºC for 10 minutes, Quant-iTTM RiboGreenTM RNA Reagent was added into well. The fluorescence intensity was measured by CLARIOstar® Plus (BMG Labtech).

#### HEK293T cell transfection

HEK293T cells were plated at 5*10^5^ cells per well in a 6-well plate in 2.5 mL DMEM media. 1 μg of the GFP mRNA was formulated with the corresponding LNP by the procedure mentioned above and then added to the cells. 24 h post-transfection, GFP expression was monitored by fluorescence microscopy.

#### Splenic cells preparation and BMDCs culture

To prepare splenic cells, mouse spleen was homogenized with the frosted end of glass slide, treated with RBC lysis buffer (Sigma) to deplete red blood cells (RBCs), followed by passing through the cell strainer (BD Biosciences). Bone-marrow derived dendritic cells (BMDCs) were prepared as described.(1) Briefly, bone marrow was isolated from mouse femurs and tibiae and treated with RBC lysis buffer (Sigma-Aldrich) to deplete RBCs. Cells were then cultured in RPMI-1640 containing 10% heat inactivated FBS (Thermo Fisher Scientific), 1% Penicillin/Streptomycin (Thermo Fisher Scientific), 50 μM 2-mercaptoethanol (Thermo Fisher Scientific), and 20 ng/ml recombinant mouse GM-CSF (eBioscience) at a density of 2 × 10^5^ cells/ml. The cells were supplemented with an equal volume of the complete culture medium described above on day 3 and refreshed with one-half the volume of medium on day 6 and harvested on day 8.

#### Incubation of LNPs with splenic cells and BMDCs

Splenic cells or BMDCs were incubated with different FITC-labeled LNP formulations (1 ug/mL) (mRNA-LNP, mRNA-BLNP1, and mRNA-BLNP2) in RPMI-1640 at 37 ℃ for 1 hours. Cells were blocked with Fc receptor binding inhibitor (clone: 93, eBioscience) for 20 minutes. Splenocytes were stained with antibodies against CD3 (clone: 17A2, BV421-conjugated, Biolegend), CD19 (clone: 1D3, PECy7-conjugated, BD Biosciences). BMDCs were stained with antibody against CD11c (clone N418 APC-conjugated, Biolegend). Labeled cells were analyzed using FACSC and Flow Cytometer (BD Biosciences).

#### Flow Cytometry

After incubation with different mRNA-LNPs, BMDC cells were washed with ice-cold FACS buffer (1% FBS in 1 × DPBS with 0.1% Sodium Azide), and incubated with purified anti-mouse CD16/32 antibody (BioLegend) in FACS buffer on ice for 20 min, followed by washing with FACS buffer. BMDCs were stained with APC anti-mouse CD11c antibody (BioLengend) at 4℃ for 30 min and washed with FACS buffer. Finally, BMDCs were stained with propidium iodide (Sigma-Aldrich). Flow cytometry was performed on FACSCanto flow cytometer (BD Bioscience).

#### Measurement of serum IgG titer

ELISA was used to determine the IgG titer of the mouse serum. The wells of a 96-well ELISA plate (Greiner Bio-One) were coated with 100 ng SARS-CoV-2 spike protein (ACROBiosystems) in 100mM sodium bicarbonate pH 8.8 at 4°C overnight. The wells were blocked with 200 ul 5% skim milk in 1X PBS at 37 °C for 1 hour and washed with 200 ul PBST (1X PBS, 0.05% Tween 20, pH 7.4) three times. Mice serum samples with 2-fold serial dilution were added into wells for an incubation at 37 °C for 2 hours and washed with 200 ul PBST six times. The wells were incubated with 100 ul HRP conjugated anti-mouse secondary antibody (1:10000, in PBS) at 37 °C for 1 hour and washed with 200 ul PBST six times. 100 μl horseradish peroxidase substrate (1-Step™ Ultra TMB-ELISA Substrate Solution) (Thermo Scientific™) was added into wells followed by 100 ul 1M H_2_SO_4_. After incubation for 30 mins, Absorbance (OD 450 nm) was measured by SpectraMax M5.

#### Pseudovirus neutralization assay

Pseudovirus was constructed by the RNAi Core Facility at Academia Sinica using a procedure similar to that described previously.(2) Briefly, the pseudotyped lentivirus carrying SARS-CoV-2 spike protein was generated by transiently transfecting HEK-293T cells with pCMV-□ R8.91, pLAS2w.Fluc.Ppuro. HEK-293T cells were seeded one day before transfection, and indicated plasmids were delivered into cells by using TransITR-LT1 transfection reagent (Mirus). The culture medium was refreshed at 16 hr and harvested at 48 hr and 72hr post-transfection. Cell debris was removed by centrifugation at 4,000 xg for 10 min, and the supernatant was passed through 0.45-□ m syringe filter (Pall Corporation). The pseudotyped lentivirus was aliquot and then stored at -80°C. To estimate the lentiviral titer by AlarmaBlue assay (Thermo Scientific), The transduction unit (TU) of SARS-CoV-2 pseudotyped lentivirus was estimated by using cell viability assay in responding to the limited dilution of lentivirus. In brief, HEK-293T cells stably expressing human ACE2 gene were plated on 96-well plate one day before lentivirus transduction. For the titering pseudotyped lentivirus, different amounts of lentivirus were added into the culture medium containing polybrene (final concentration 8 □ g/ml). Spin infection was carried out at 1,100 xg in 96-well plate for 30 minutes at 37 °C. After incubating cells at 37°C for 16 hr, the culture medium containing virus and polybrene were removed and replaced with fresh complete DMEM containing 2.5 μg/ml puromycin. After treating puromycin for 48 hrs, the culture media was removed, and the cell viability was detected by using 10% AlamarBlue reagents according to manufacturer’s instruction. The survival rate of uninfected cells (without puromycin treatment) was set as 100%. The virus titer (transduction units) was determined by plotting the survival cells versus diluted viral dose. For neutralization assay, heat-inactivated sera or antibodies were serially diluted and incubated with 1,000 TU of SARS-CoV-2 pseudotyped lentivirus in DMEM for 1 h at 37°C. The mixture was then inoculated with 10,000 HEK-293T cells stably expressing human ACE2 gene in a 96-well plate. The culture medium was replaced with fresh complete DMEM (supplemented with 10% FBS and 100 U/mL penicillin/streptomycin) at 16 h postinfection and continuously cultured for another 48 h. The expression level of luciferase gene was determined by using Bright-Glo Luciferase Assay System (Promega). The relative light unit (RLU) was detected by Tecan i-control (Infinite 500). The percentage of inhibition was calculated as the ratio of RLU reduction in the presence of diluted serum to the RLU value of no serum control using the formula (RLU^control^ - RLU^Serum^)/RLU control.

#### FACS analysis of cell-type specific transfection in lymph nodes

To assess the in vivo cellular targeting of eGFP-mRNA LNPs, seven mice were analyzed, including six treated animals (n = 3 per group for two different GFP-mRNA LNP formulations) and one untreated control (n = 1). After 6 hours post-injection, inguinal lymph nodes (iLNs) were harvested for analysis. Cell suspensions from iLNs were stained using a viability dye (FVS440UV) followed by antibodies targeting CD3 (T cells), CD19 (B cells), CD11c (dendritic cells), F4/80 (macrophages), CD80, and CD86 (maturation markers). eGFP expression was used to identify successfully transfected cells. A total of 14 FACS tubes were analyzed, enabling quantification of dendritic cells (CD11c^+^), mature dendritic cells (CD11c^+^CD80^+^ and CD11c^+^CD86^+^), macrophages (F4/80^+^), B cells (CD19^+^), and T cells (CD3^+^).

#### Flow cytometric gating strategy and identification of CD3+CD8+Granzyme B (Grz B) producing T cells

The inguinal draining lymph node cells of BALB/c mice immunized with influenza mRNA vaccine or PBS were harvested on day 14 following second dose immunization. All cells obtained from each immunized mouse were re-stimulated with H1-specific HA (H1N1/Victoria/2019) peptide pools for 48 hours at 37°C. Singlets were gated by FSC and SSC (P1∼P3). Live cells were distinguished by Live/Dead Red dye (P4). Next, CD3+CD8+ T cell population was further selected in P5. Finally, Granzyme B-producing T cells within CD3+CD8+T cell population were then identified (P6).

#### Analysis of cytotoxic T cell immunity induced by influenza mRNA vaccines in mice Five

BALB/c mice per group were intramuscularly immunized with either the indicated influenza mRNA vaccine or PBS. Inguinal draining lymph nodes (dLNs) were harvested, and cells were re-stimulated with H1 HA peptide pool (H1N1/Victoria/2019) for 48 hours at 37°C. Brefeldin A, a protein transport inhibitor, was added at the last 4 hours of culture. Subsequently, the cells were harvested and stained with T cell surface markers (CD3 and CD8), followed by fixed, permeabilized, and intracellularly stained with granzyme B (GrzB) marker. (A) The percentage of GrzB-producing cells in CD3+CD8+ T cell populations and (B) the numbers of CD3+CD8+GrzB-producing cell in inguinal dLNs were further quantified by flow cytometric analysis. Mean ± standard error of the mean (SEM) was plotted, and the statistical significances compared to PBS group were determined by one-way ANOVA with Prism 10.4.1. *p < 0.05; ***p < 0.001; ****p < 0.0001.

### Synthesis

**Scheme S1.**
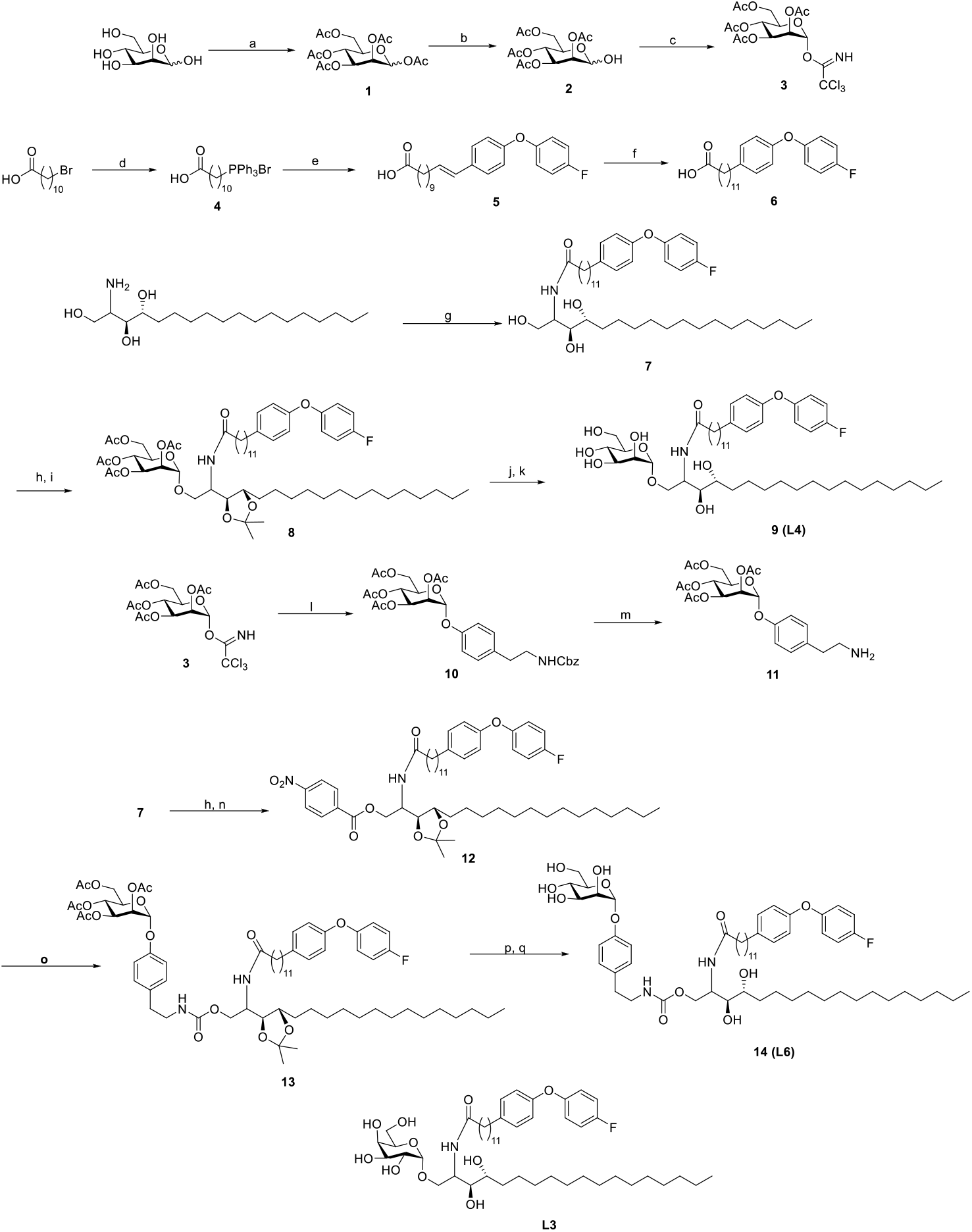
Synthetic scheme of compound **L3, L4** and **L6**. Reagents and conditions were as follows. (a)Ac_2_O, pyridine, 2h, quant.(b) hydrazine acetate, DMF, 1h, 40°C, 85% (c) trichloroacetonitrile, DBU, DCM, 4h, 82% (d) PPh_3_, toluene, reflux, 24h, 76% (e) 4-(4-fluorophenoxy)benzaldehyde, LHMDS, THF, 0°C to rt, 12h, 69% (f) H_2_, Pd/C, MeOH, 2h, quant.(g) **6**, HBTU, DIPEA, DCM, 12h, 80% (h) 2,2-DMP, PTSA, 2h, quant. (i) BF_3_OEt_2_, 4A MS, DCM, 12h, 71% (j) NaOMe, MeOH, 2h, quant. (k) TFA, DCM, 12h, 79% (l) BF_3_OEt_2_, 4A MS, DCM, 12h, 82% (m) H_2_, Pd/C, AcOH, MeOH, 1h. (n) 4-nitrophenyl chloroformate, Et_3_N, THF, 12h (o) **11**, Et_3_N, DMF, 12h, 2 steps 34% (p) NaOMe, MeOH, 2h, quant. (q) TFA, DCM, 12h, 71%

**Scheme S2.**
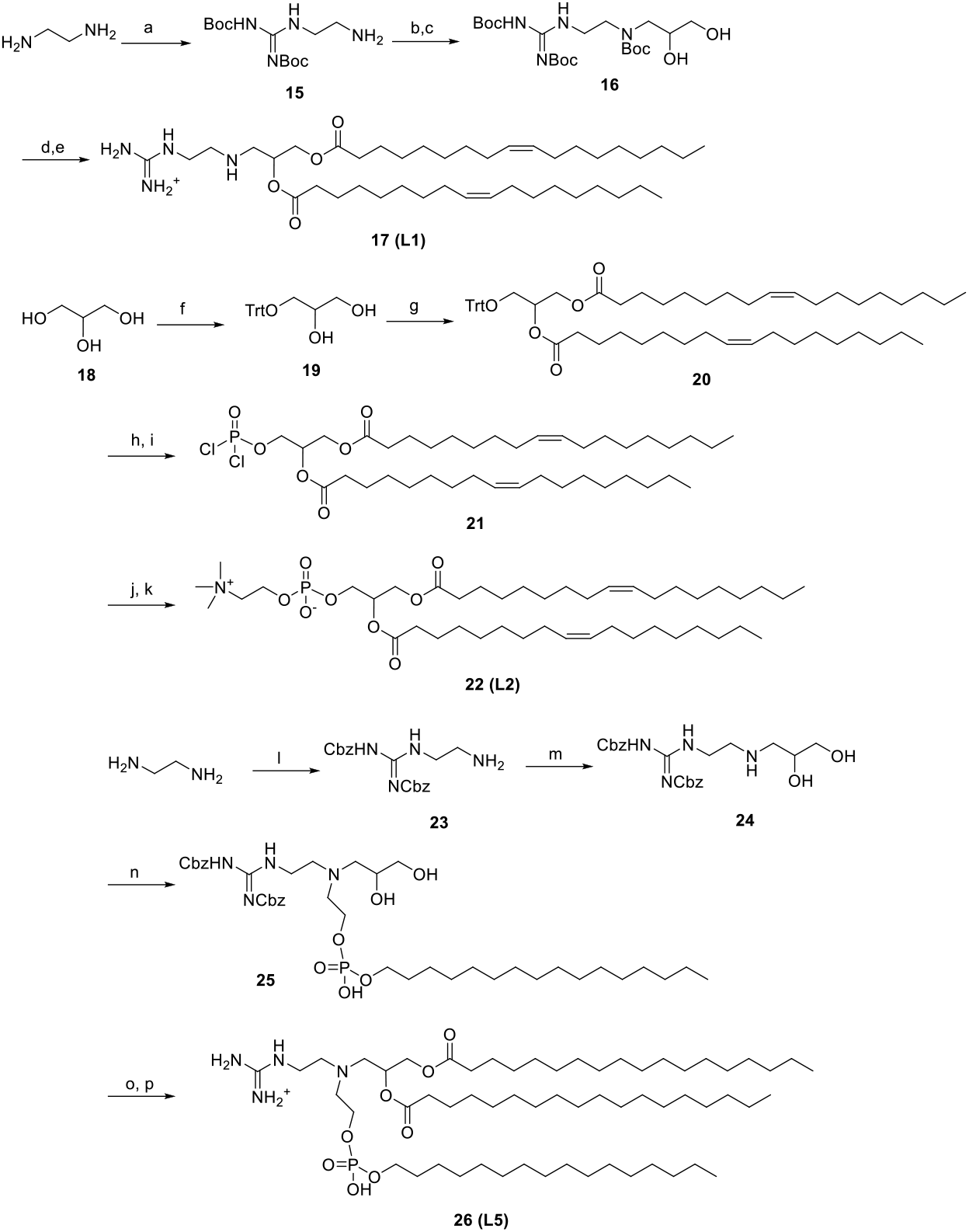
Synthetic procedure of **L1, L2**, and **L5**. Reagents and conditions were as follows. (a) N,N-di-Boc-1H-pyrazole-1-carboxamidine, DIPEA, DCM, 2h, 90%. (b) 3-bromo-1,2-propanediol, K_2_CO_3_, MeCN, 95°C, 3h, 32%. (c) Boc_2_O, DCM, 2h, quant.. (d) oleic acid, DMAP, EDC, Et_3_N, DCM, 2h, 67%. (e) TFA, DCM, 1h, quant. (f) TrtCl, DMAP, THF, 12h, 85%. (g) oleic acid, DMAP, EDC, DCM, 12h, 70%. (h) HCl, DCM, MeOH, 4°C, 12h, 60% (i) POCl_3_, pyridine, toluene, 0 °C, 4h. (j) choline tosylate, CHCl3, pyridine, 12h. (k) 10 % aq. NaHCO_3_, 1 h, 3 steps 40%. (l) N,N-bis(benzyloxycarbonyl)-1H-pyrazole-1-carboxamidine, DIPEA, DCM, 2h, 88%. (m) 3-bromo-1,2-propanediol, K_2_CO_3_, MeCN, 95°C, 3h, 35%. (n) 2-(hexadecyloxy)-1,3,2-dioxaphospholane 2-oxide, DMF, 70°C, 24h, 70%. (o) stearic acid, EDC, DMAP, DCM, 12h, 65%. (p) H_2_, Pd/C, MeOH, 12h, quant.

Compound **3 (3), 6 (4), 11 (5), L3 (4), 20 (6)** were synthesized according to published articles.

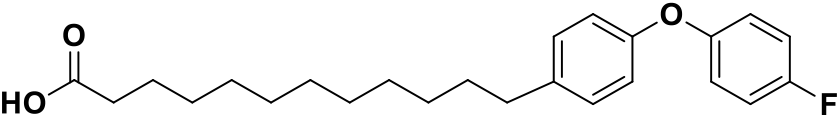

**Compound 6**. (11-Carboxynonyl)triphenylphosphonium bromide **4** (2.5 g, 10 mmol) was prepared by refluxing triphenylphosphine (10 mmol) and 11-bromoundecanoic acid (10 mmol). It was then dissolved in 50 ml of tetrahydrofuran (THF) and cooled to 0 °C. lithium bis(trimethylsilyl)amide (LHMDS; 1 M in THF, 20 mmol) was added to the solution to produce an orange ylide. After that, 4-(4-Fluorophenoxy)benzaldehyde (12 mmol) in 20 ml of THF was added dropwise to the solution and stirred for 4 h at room temperature. The reaction was quenched with methanol and concentrated. The residue was extracted with EA and brine and then dried over MgSO_4_. After removal of the solvent, the mixture was chromatographed on silica gel (EA-Hex =1:2) to give the unsaturated fatty acid **5**. The saturated fatty acid was prepared by catalytic hydrogenation in 50 ml of methanol containing 10 mol% of 10% palladium on charcoal (Pd/C). The reaction mixture was stirred under H_2_ at room temperature overnight. The hydrogenated product was filtered through Celite, and the resulting solution was concentrated to give the product (1.5 g, 66%). ^1^H NMR (600 MHz, CDCl_3_) δ 7.15 (m, 2H), 7.07 (m, 2H), 6.96 (m, 2H), 6.87 (m, 2H), 2.58 (t, J = 7.6 Hz, 2H), 2.26 (m, J = 7.6 Hz, 2H), 1.59 (m, 4H), 1.32 (m, 14H).

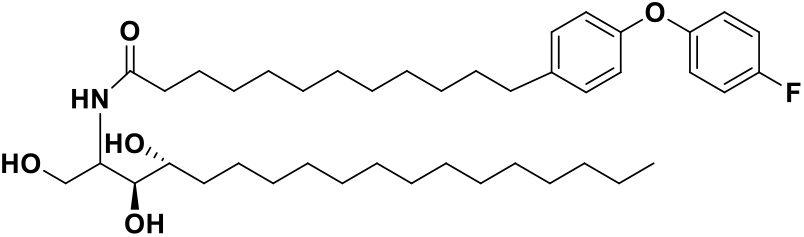

Compound **7**. Compound **6** (1 mmol) in THF (10 mL) was added EDC (1.5 mmol), HOBt (1.5 mmol), DMAP (0.1 mmol), trimethylamine (2 mmol), and phytosphingosine (1.2 mmol), and the resulting solution was stirred under nitrogen at rt for 12 h. The solvent was then removed by evaporation, followed by extraction with EA/H_2_O. The collected organic layer was washed with saturated NaHCO_3_(aq), water and brine, and dried over MgSO_4_. The crude product was purified by column chromatography on silica gel (EA/Hex 1:1) to yield **7** (0.5 g, 74%). ^1^H NMR (600 MHz, CDCl_3_) δ 7.28 (d, J = 8.4 Hz, 2H), 7.15 (t, J = 8.4 Hz, 2H), 7.04-7.09 (m, 4H), 5.59 (d, J = 3.8 Hz, 1H), 4.45-4.38 (m, 2H), 4.35-4.30 (m, 1H), 2.59 (t, J = 7.6 Hz, 2H), 2.45 (t, 2H), 2.30 (m, 1H), 1.91 (m, 2H), 1.81 (m, 2H), 1.68 (m, 2H), 1.59 (m, 2H), 1.48-1.15 (m, 40H), 0.87 (t, J = 6.9 Hz, 3H).

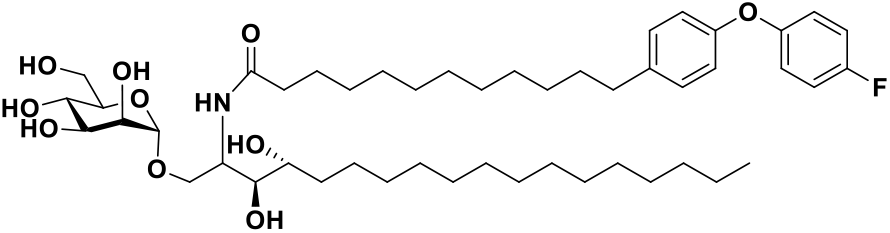

**Compound 9**. Compound **7** (1 mmol) in 2,2-DMP (10 mL) was added PTSA (0.03 mmol) and the resulting mixture was stirred at rt for 2h. The mixture was then dried by evaporation, followed by extraction with EA/H_2_O. The collected organic layer was washed with water and brine and dried over MgSO_4_. The crude product was purified by column chromatography on silica gel (EA/Hex 1:1). After that, to a stirred solution of purified compound and 4 A molecular sieve (0.1 g) in anhydrous DCM (5 mL) was cooled to -40°C and then BF_3_.OEt_2_ (0.1 mmol) was added dropwise to the solution. A solution of **3** in anhydrous DCM was added dropwise to the above mixture and stirred for 1 h at -40°C. After that, the reaction gradually warmed to room temperature and stirred for another 1 h. The solution was quenched by adding triethylamine, then filtered, added sat. NaHCO_3_ aq. and extracted with DCM. The organic layer was dried with MgSO_4_ and evaporated to dryness. The residue was purified by flash column chromatography on silica gel to give **8**. The product was then dissolved in MeOH and NaOMe (0.2 eq) was added, and the resulting solution was stirred at rt for 2 h. The mixture was neutralized by IR-120, and then filtered, concentrated to dryness in vacuo.in MeOH was added NaOMe (0.2 eq) and the resulting solution was stirred under nitrogen at rt for 2 h. The mixture was neutralized by IR-120, and then filtered, concentrated to dryness in vacuo. Finally, the product was treated with 10% TFA in DCM to give the deprotected compound **9** (0.47 g, 4 steps 56%). ^1^H NMR (600 MHz, CDCl_3_:MeOD=1:1) δ 7.12-7.10 (d, *J* = 8.4 Hz, 2H), 7.01-6.92 (m, 4H), 6.86-6.84 (d, *J* = 8.4 Hz, 2H), 4.77 (s, 1H), 4.17-4.15 (m, 1H), 3.87-3.49 (m, 10H), 2.56 (t, *J* = 7.5 Hz, 2H), 2.20 (t, *J* = 7.5 Hz, 2H), 1.62-1.52 (m, 5H), 1.30-1.24 (m, 39H), 0.86 (t, *J* = 6.9 Hz, 3H). ^13^C NMR (150 MHz, CDCl_3_:MeOD=1:1) δ 175.63, 160.21, 156.16, 154.30, 138.70, 130,32(x2), 120.74, 120.69, 119.08 (x2), 116.86, 116.70, 101.45, 75.09, 73.80, 72.81, 71.96, 71.39, 67.95, 67.53, 62.18, 51.86, 51.77, 37.11, 37.06, 35.88, 32.70, 32.65, 32.39, 30.49, 30.44 (x6), 30.38, 30.34, 30.28, 30.25, 30.18, 30.08, 29.99,. 26.72, 26.63, 23.36, 14.52. HRMS (ESI) calcd for C_48_H_79_FNO_10_^+^ [M + H]^+^: 848.5688, found 848.5676.

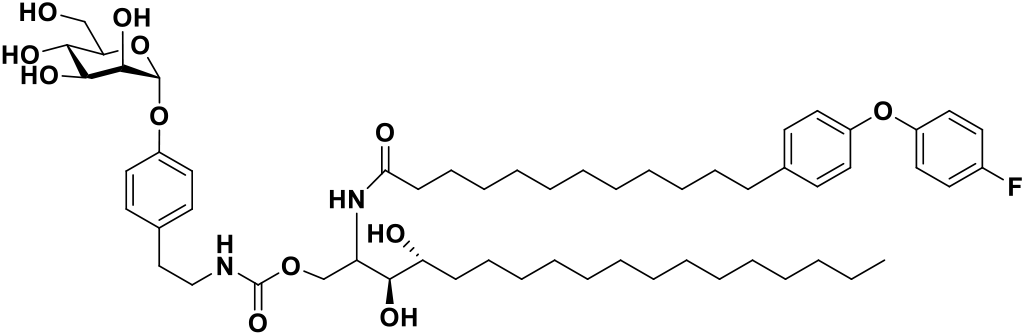

Compound **14**. Compound **7** (1 mmol) in 2,2-DMP (10 mL) was added PTSA (0.03 mmol) and the resulting mixture was stirred at rt for 2h. The mixture was then dried by evaporation, followed by extraction with EA/H_2_O. The collected organic layer was washed with water and brine and dried over MgSO_4_. The crude product was purified by column chromatography on silica gel (EA/Hex 1:1). Purified compound in THF (10 mL) was added 4-nitrophenylchloroformate (2 mmol), trimethylamine (2 mmol), and the resulting solution was stirred under nitrogen at rt for 12 h. The solvent was then removed by evaporation, and the crude compound was directly used for next step without further purification. Compound **11** (1.5 mmol) in THF (10 mL) was added **12** (1 mmol) and trimethylamine (2 mmol), and the resulting solution was stirred under nitrogen at rt for 2 h. The solvent was then removed by evaporation, followed by extraction with EA/H_2_O. The collected organic layer was washed with saturated NaHCO_3_(aq), water and brine, and dried over MgSO_4_. The crude product was purified by column chromatography on silica gel (EA/Hex 1:1+10% MeOH) to yield **13** (34%). Compound **13** in MeOH was added NaOMe (0.2 eq) and the resulting solution was stirred under nitrogen at rt for 2 h. The mixture was neutralized by IR-120, and then filtered, concentrated to dryness in vacuo. Finally, the product was treated with 10% TFA in DCM to give the fully deprotection compound **14** (0.2 g, 2 steps 71%). ^1^H NMR (600 MHz, CDCl_3_/MeOD=1:1) δ 7.16-7.13 (m, 4H), 7.05-7.00 (m, 4H), 6.95-6.92 (m, 2H), 6.87 (d, J = 8.6 Hz, 2H), 5.49 (s, 1H), 5.3 (s, 1H), 4.41-4.32 (m, 2H), 4.01 (s, 1H), 3.96-3.94 (m, 1H), 3.81-3.70 (m, 3H), 3.65-3.60 (m, 1H), 3.51-3.48 (m, 2H), 3.40-3.35 (m, 2H), 2.71 (t, J = 7.6 Hz, 2H), 2.55 (t, J = 7.6 Hz, 2H), 2.30 (t, J = 7.6 Hz, 2H), 1.65-1.58 (m, 7H), 1.42-1.25 (m, 42H), 0.87 (t, J = 6.9 Hz, 3H). ^13^C NMR (150 MHz, CDCl_3_/MeOD=1:1) δ 174.71, 159.37, 157.5, 155.31, 154.95, 153.47, 137.86, 132.77, 129.46, 119.88, 119.82, 118.22, 116.51, 115.99, 115.84, 98.61, 77.72, 77.51, 77.29, 74.38, 73.31, 71.81, 71.09, 70.56, 66.85, 63.71, 61.22, 50.37, 48.70, 48.56, 48.41, 48.27, 48.13, 47.99, 47.84, 42.26, 36.30, 35.11, 35.00, 32.07, 31.78, 31.51, 29.55, 29.50, 29.44, 29.37, 29.35, 29.29, 29.20, 29.15, 29.09, 25.77, 25.68, 22.49, 13.60. HRMS (ESI) calcd for C_57_H_88_FN_2_O_12_^+^ [M + H]^+^: 1011.6321, found 1011.6349.

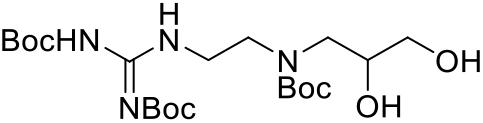

Compound **16**. Ethylenediamine (100 mmol) in DCM (50 ml) was added a solution of N,N-di-Boc-1H-pyrazole-1-carboxamidine (10 mmol) and DIPEA (12 mmol) in DCM (50 ml) at 0°C, and the resulting mixture was warmed to rt and stirred for 2h. The mixture was then dried by evaporation, followed by extraction with EA/H_2_O. The collected organic layer was washed with water and brine and dried over MgSO_4_ to give compound **15**. A solution of 3-bromo-1,2-propanediol (1 eq), potassium carbonate (1.2 eq), **15** (10 eq) in MeCN (50 mL) was stirring under reflux for 3 h. The reaction mixture was cooled to rt and the solvent was removed under reduced pressure. Then water (40 mL) was added, and the resulting mixture was extracted by EA (40 mL) three times. The organic phase was combined, dried over anhydrous MgSO_4_, filtered and dried to give the crude product. The crude product was then added to a solution of Boc_2_O in DCM and stirred at rt for 2 h. The solvent was removed under reduced pressure, and the crude product was purified by column chromatography to give compound **16** (0.9 g, 32%). ^1^H NMR (600 MHz, CDCl_3_) δ 4.22 (m, 1H), 3.82 (d, *J* = 3.9 Hz, 2H), 3.59-3.53 (d, *J* = 3.9 Hz, 2H), 3.21-3.12 (m, 4H), 1.42 (m, 27H).

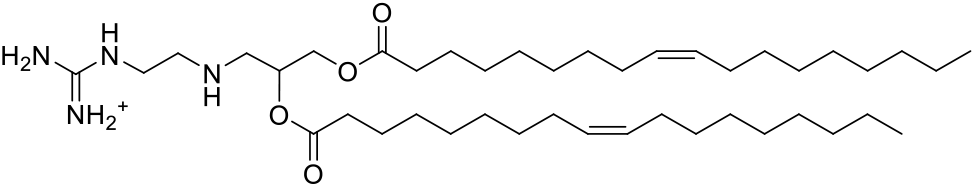

Compound **17**. To a stirred solution of **16** (0.9 mmol) in 9 mL of DCM was added oleic acid (2 mmol) and DMAP (0.2 mmol), EDC (2 mmol), and DIPEA (2 mmol). The resulting solution was stirred overnight at room temperature. After completion of reaction, the resulting mixture was extracted by DCM (40 mL) three times. The organic phase was combined, dried over anhydrous MgSO_4_, filtered and dried to give the crude product. The crude was purified by column chromatography (EA/Hex = 1:2), and the product was treated with 10% TFA in DCM to give compound **17** (0.4 g, 67%). ^1^H NMR (600 MHz, MeOD) δ 5.35-5.30 (m, 3H), 5.25-5.23 (m, 1H), 4.45-4.42 (m,1H), 4.27 (d, *J* = 3.9 Hz, 2H), 4.19-4.16 (m, 1H), 4.04 (t, *J* = 3.9 Hz, 2H), 3.63 (t, *J* = 3.9 Hz, 2H), 2.36-2.31 (m, 4H), 2.04-2.03 (m, 7H), 1.61-1.60 (m, 4H), 1.33-1.30 (m, 30H), 0.90 (t, *J* = 6.9 Hz, 6H). ^13^C NMR (150 MHz, MeOD) δ 175.12, 174.86, 158.0, 131.01 (x2), 130.71 (x2), 72.08, 71.92, 67.53, 65.01, 63.85, 60.60, 35.28, 35.07, 33.33, 31.01 (x2), 30.78 (x5), 30.61 (x5), 30.52 (x5), 30.35, 28.32, 26.18, 23.90 (x2), 14.63 (x2).

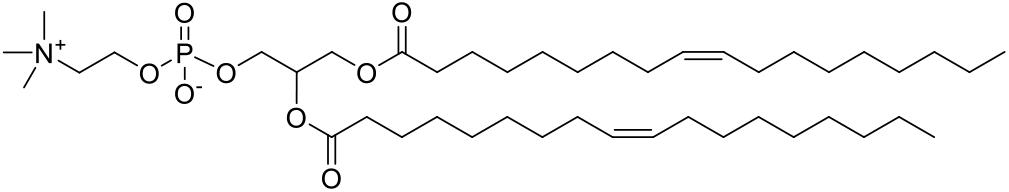

Compound **22. 20 (6)** (3 mmol) in a mixture of toluene (21 mL) and pyridine (12 mmol) were cooled to 0 °C under nitrogen atmosphere. A solution of phosphorus oxychloride (6 mmol) in toluene (9 mL) was added dropwise to the reaction mixture over 20 minutes, while keeping the temperature of the reaction mixture at 0 °C. The reaction mixture was stirred for 15 minutes, then allowed to warm to rt and stirred for an additional 4 h. The reaction mixture was evaporated to obtain crude product **21**. A solution of crude **21** in anhydrous DCM (30 mL) was cooled to 0 °C. Choline tosylate (2.7 mmol) was taken in a mixture of anhydrous DCM (30 mL) and pyridine (6 mL) and sonicated at 40 °C to obtain a clear solution. This solution was added dropwise to the solution of crude compound **21** over 20 minutes, while maintaining the temperature at 0 °C. The reaction mixture was stirred for 20 minutes, then allowed to warm to rt and stirred for additional 18 h. The reaction mixture was quenched by addition of 10% NaHCO_3(aq)_ solution and diluted with DCM (50 mL). The aqueous layer was extracted with DCM, and the combined organic layers were dried over MgSO_4_. Solvent was evaporated and the crude product was purified by silica gel column chromatography using DCM:MeOH:H_2_O system to afford **22** (0.94 g, 40%). ^1^H NMR (600 MHz, MeOD) δ 5.37-5.32 (m, 3H), 5.26-5.24 (m, 1H), 4.45-4.43 (m,1H), 4.28 (d, *J* = 3.9 Hz, 2H), 4.19-4.16 (m, 1H), 4.00 (t, *J* = 3.9 Hz, 2H), 3.64 (t, *J* = 3.9 Hz, 2H), 3.31 (s, 9H), 2.36-2.31 (m, 4H), 2.04-2.03 (m, 7H), 1.61-1.60 (m, 4H), 1.33-1.30 (m, 30H), 0.90 (t, *J* = 6.9 Hz, 6H). ^13^C NMR (150 MHz, MeOD) δ 175.03, 174.73, 131.06 (x2), 130.91 (x2), 71,98, 71.92, 67.63, 65.01, 63.85, 60.60, 54.84 (x3), 35.24, 35.07, 33.23, 31.01 (x2), 30.78 (x5), 30.62 (x5), 30.52 (x5), 30.37, 28.32, 26.16, 23.90 (x2), 14.63 (x2).

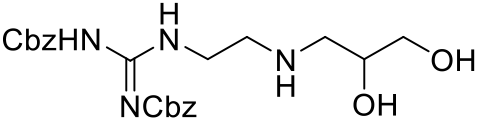

Compound **24**. Ethylenediamine (100 mmol) in DCM (50 ml) was added to the solution of N,N-bis(benzyloxycarbonyl)-1H-pyrazole-1-carboxamidine (10 mmol) and DIPEA (12 mmol) in DCM (50 ml) at 0 °C, and the resulting mixture was warmed to rt and stirred for 2h. The mixture was then dried by evaporation, followed by extraction with EA/H_2_O. The collected organic layer was washed with water and brine and dried over MgSO_4_ to give compound **23**. A solution of 3-bromo-1,2-propanediol (1 eq), potassium carbonate (1.2 eq), **23** (10 eq) in MeCN (50 mL) was stirring under reflux for 3 h. The reaction mixture was cooled to rt and the solvent was removed under reduced pressure. Then water (40 mL) was added, and the resulted mixture was extracted by EA (40 mL) for three times. The organic phase was combined, dried over anhydrous MgSO_4_, filtered and dried. The crude product was purified by silica gel column chromatography to give the compound **24** (0.9 g, 35%). 1H NMR (600 MHz, MeOD): □ 11.52 (br, 1H), 8.62 (s, 1H), 7.41-7.29 (m, 11H), 5.16 (s, 2H), 4.19 (m, 1H), 3.80 (d, *J* = 3.9 Hz, 2H), 3.58-3.52 (d, *J* = 3.9 Hz, 2H), 3.45 (t, *J* = 3.9 Hz, 2H), 2.86 (t, *J* = 3.9 Hz, 2H).

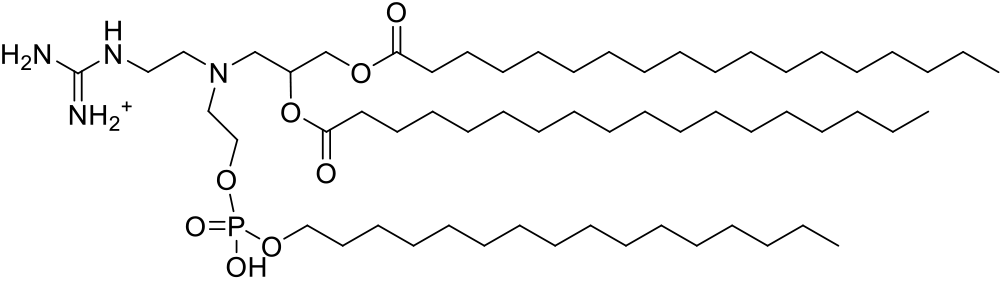

Compound **26**. A solution of **24** (0.25 mmol) in 1 ml anhydrous DMF was added 2-(hexadecyloxy)-1,3,2-dioxaphospholane 2-oxide (7) (0.25 mmol), and the reaction was stirred at 70 °C for 24 h. The mixture was concentrated to dryness *in vacuo* to give **25**. To a stirred solution of **25** (0.2 mmol) in 2 mL of DCM was added stearic acid (0.44 mmol) and DMAP (0.2 mmol), EDC (0.44 mmol), and DIPEA (0.66 mmol). The resulting solution was stirred overnight at room temperature. After completion of reaction, the resulting mixture was extracted by DCM (40 mL) three times. The organic phase was combined, dried over anhydrous MgSO_4_, filtered and dried to give the crude product. The crude was purified by column chromatography (EA/Hex = 1:4), and the product was prepared by catalytic hydrogenation in 2 ml of methanol containing 10 mol% of 10% palladium on charcoal (Pd/C). The reaction mixture was stirred under H_2_ at room temperature overnight. The hydrogenated product was filtered through Celite, and the resulting solution was concentrated to give **26** (0.11 g, 3 steps 45%). ^1^H NMR (600 MHz, MeOD) δ 4.50-4.38 (m, 4H), 4.45-4.42 (m,1H), 4.27 (d, *J* = 3.9 Hz, 2H), 4.19-4.16 (m, 3H), 4.04 (t, *J* = 3.9 Hz, 2H), 3.63 (t, *J* = 3.9 Hz, 2H), 2.36-2.31 (m, 4H), 2.04-2.03 (m, 7H), 1.74-1.70 (m, 2H), 1.61-1.60 (m, 4H), 1.38-1.20 (m, 58H), 0.90 (t, *J* = 6.9 Hz, 9H).

**Figure S1.**
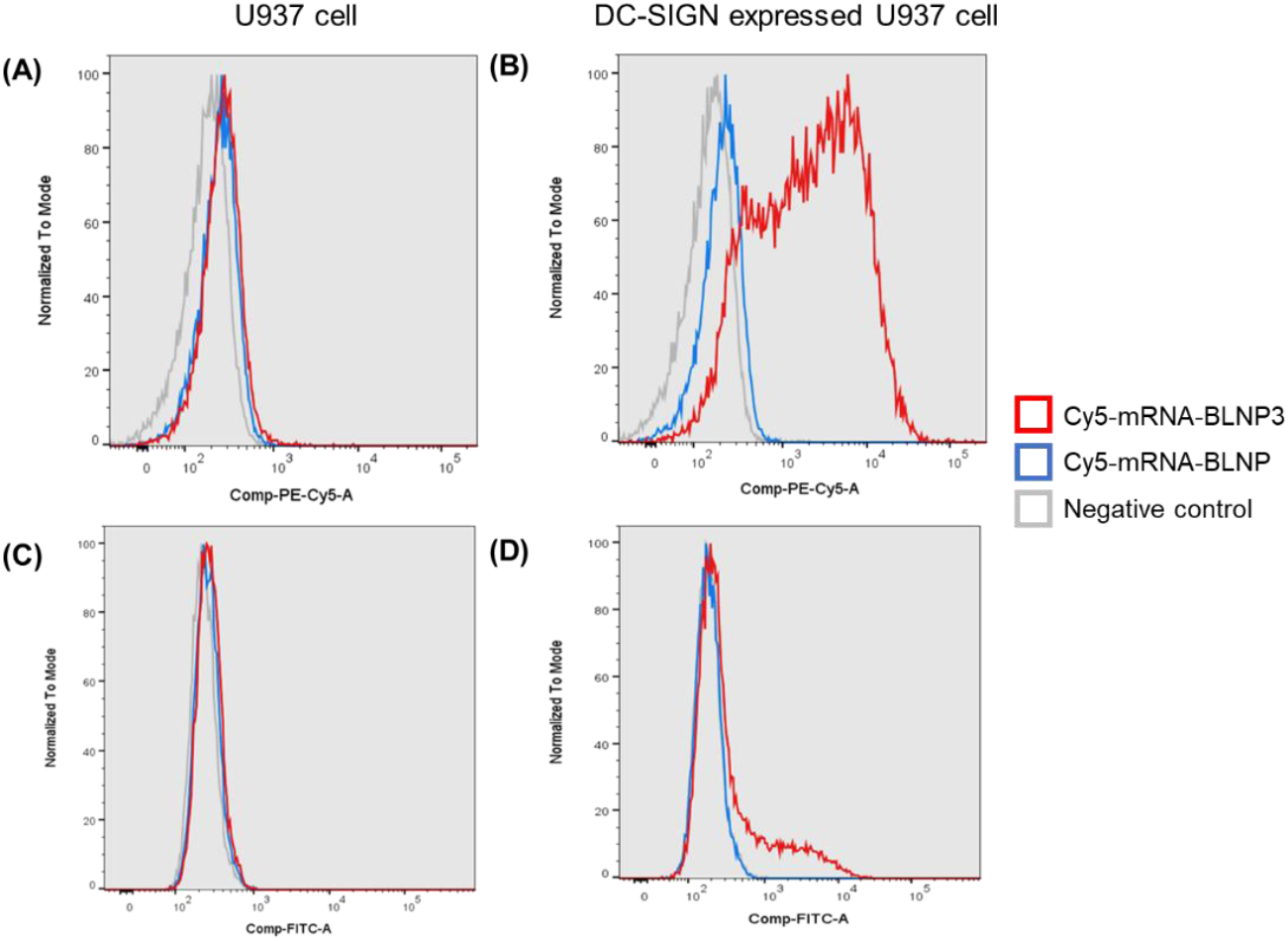
Cellular uptake of mRNA-LNPs by (A) U937 cell and (B) DC-SIGN expressed U937 cell and transfection by (C) U937 cell and (D) DC-SIGN expressed U937 cell.

**Figure S2.**
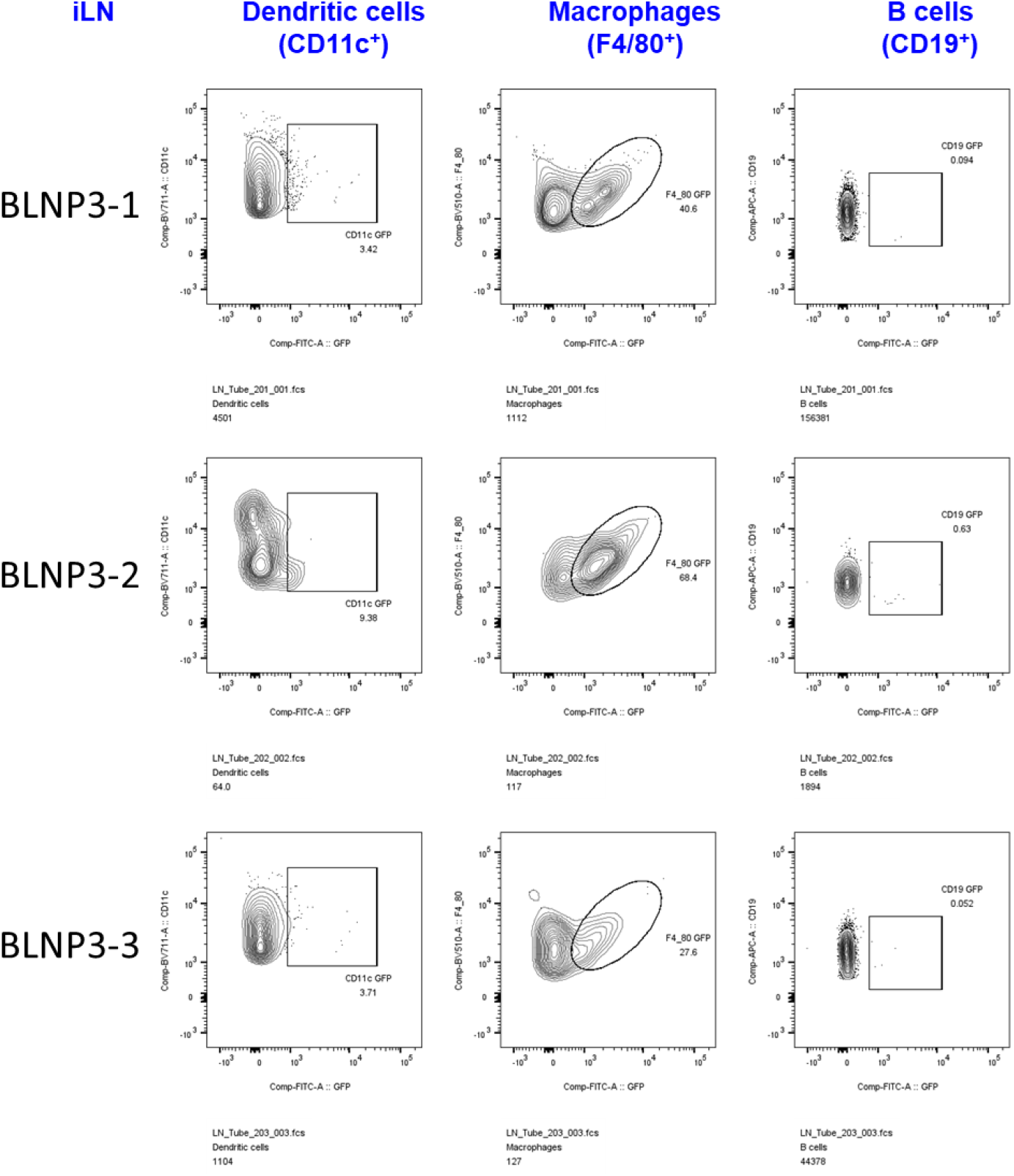
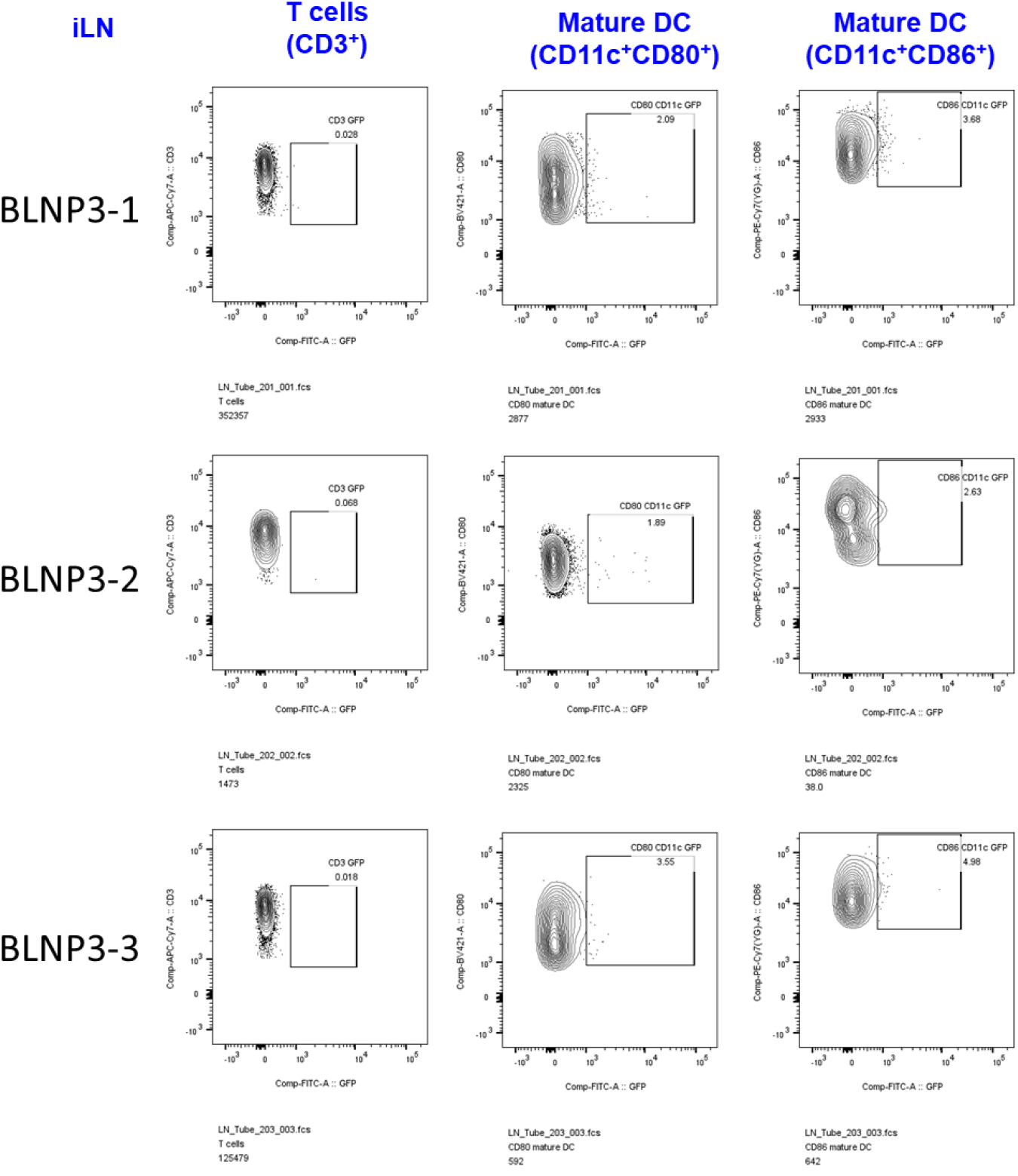
Representative diagrams of flow cytometry for analysis of eGFP positive cells with LNs after treatment with eGFP-mRNA BLNP3.

**Figure S3.**
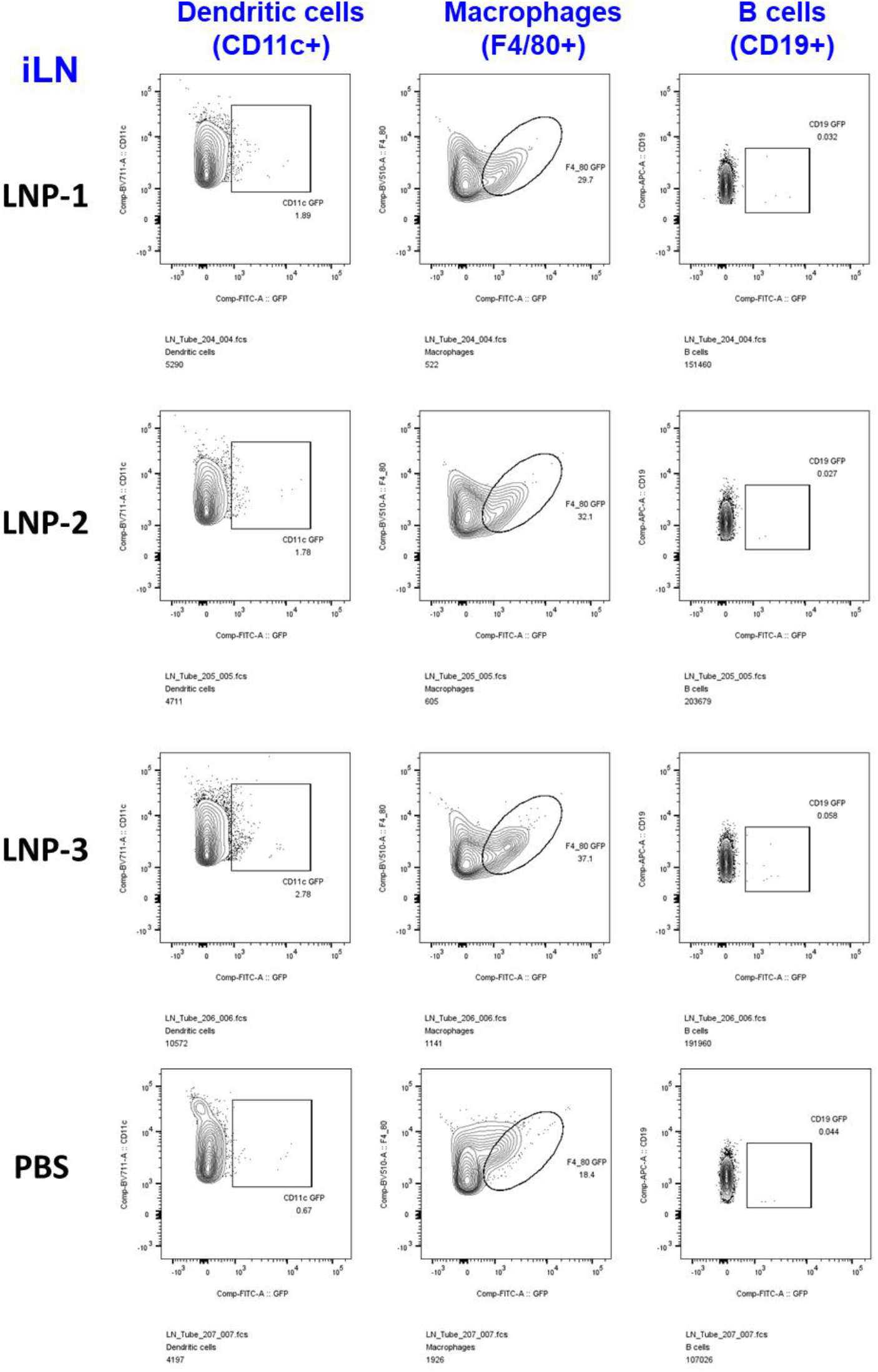
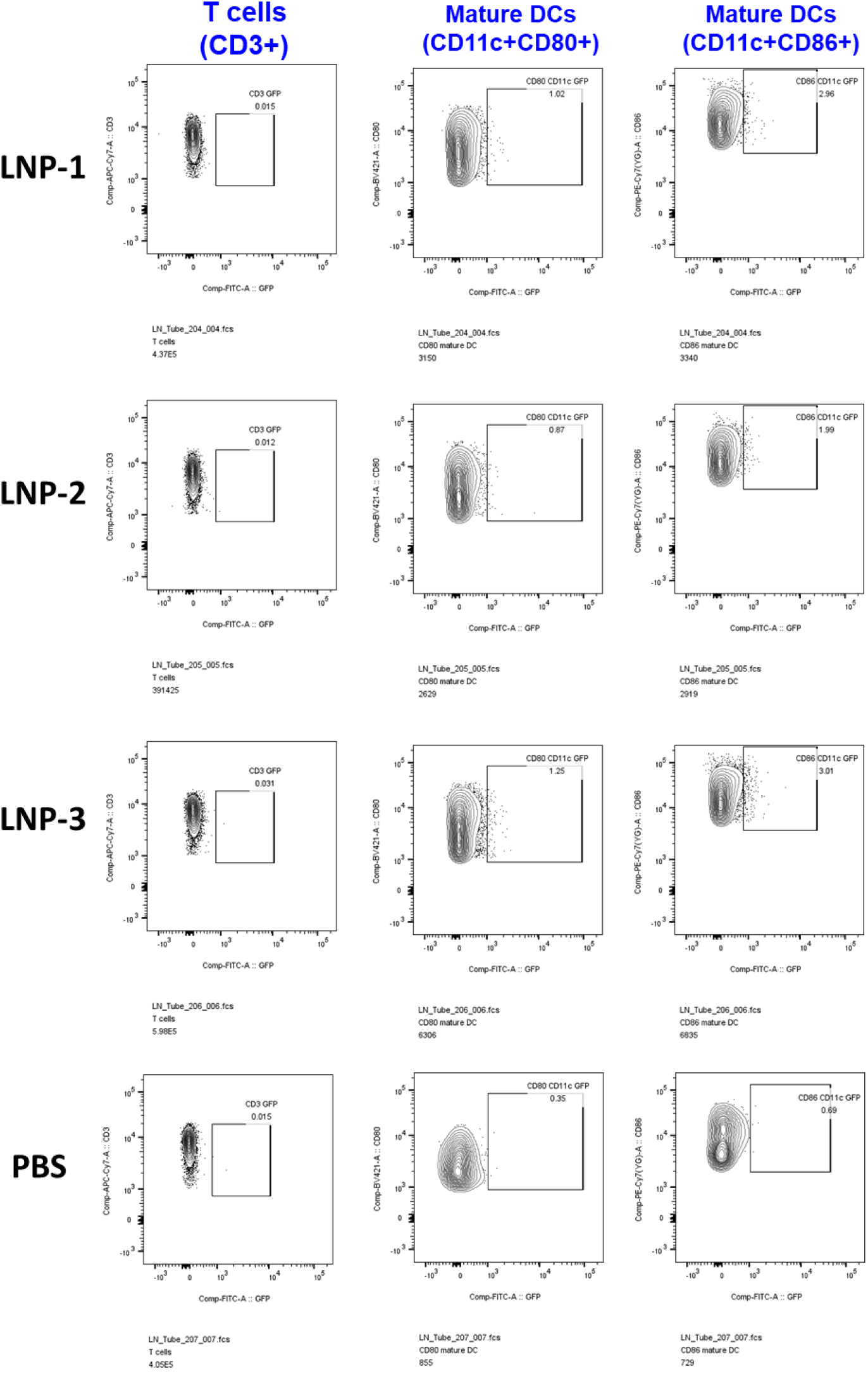
Representative diagrams of flow cytometry for analysis of eGFP positive cells with LNs after treatment with eGFP-mRNA LNP and PBS.

**Figure S4.**
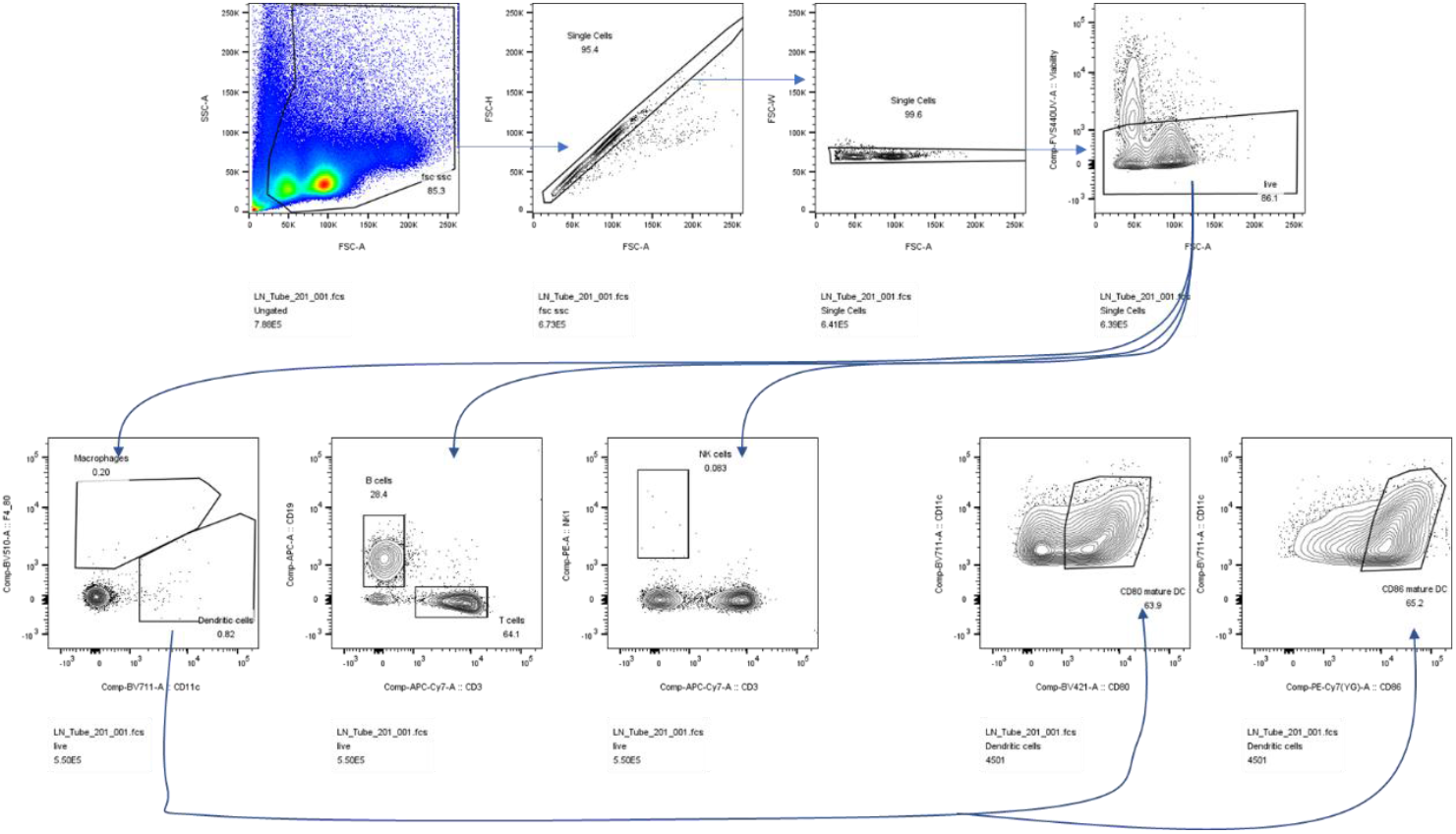
Strategy for iLN cell gating in flow cytometry.

**Figure S5.**
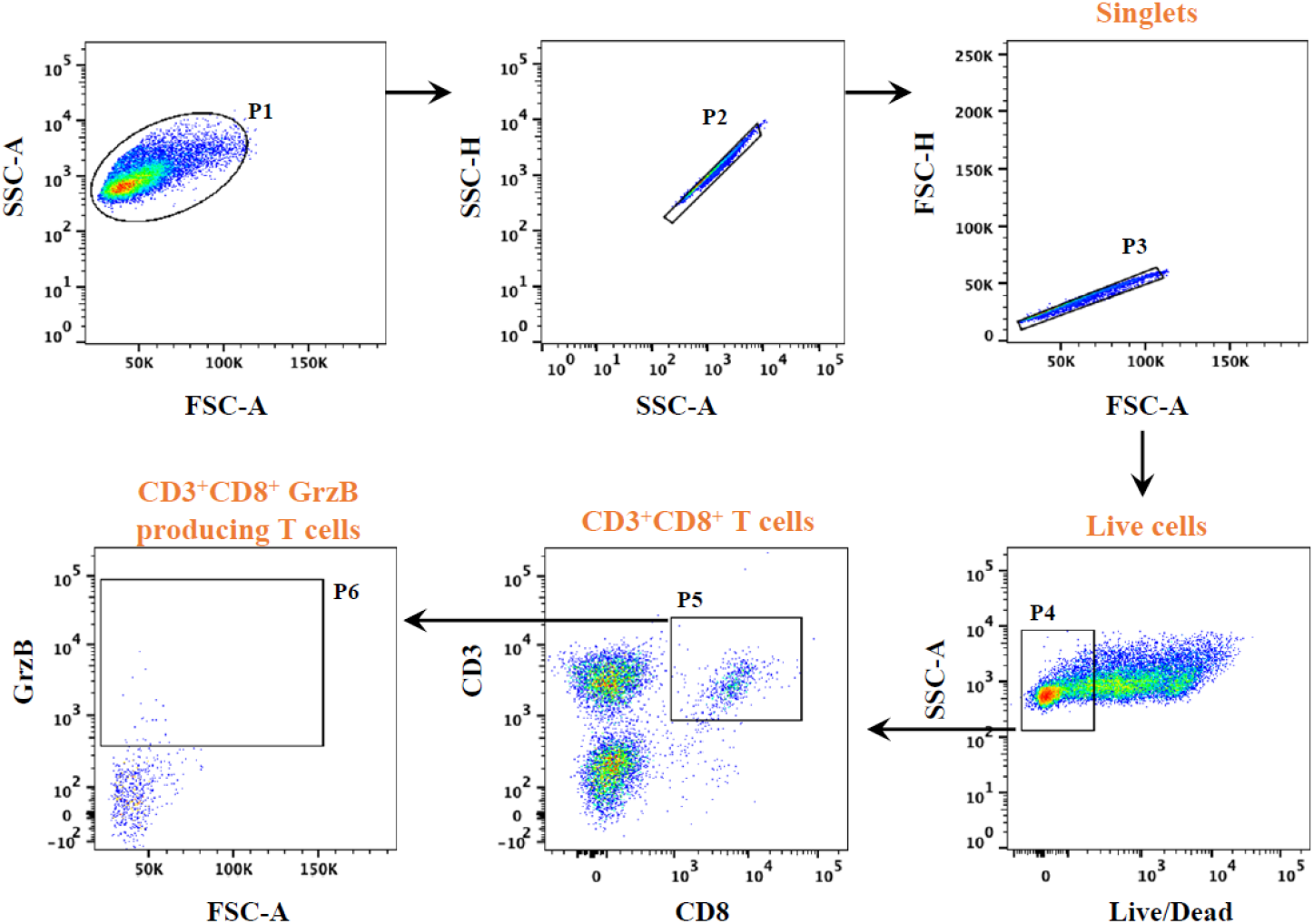
Flow cytometric gating strategy and identification of CD3+CD8+Granzyme B (Grz B) producing T cells. The inguinal draining lymph node cells of BALB/c mice immunized with influenza mRNA vaccine or PBS were harvested on day 14 following second dose immunization. All cells obtained from each immunized mouse were re-stimulated with H1-specific HA (H1N1/Victoria/2019) peptide pools for 48 hours at 37°C. Singlets were gated by FSC and SSC (P1∼P3). Live cells were distinguished by Live/Dead Red dye (P4). Next, CD3+CD8+ T cell population was further selected in P5. Finally, Granzyme B-producing T cells within CD3+CD8+T cell population were then identified (P6).

